# Secreted ORF8 reprograms macrophages to enhance SARS-CoV-2 infection of lung epithelial cells

**DOI:** 10.64898/2026.06.10.731427

**Authors:** Yusuke Matsui, Rahul K. Suryawanshi, Mauricio Montano, Taha Y. Taha, Mir M. Khalid, Limeng Sun, Kanika Khanna, Jin Tang, Yuan Zhou, Robyn M. Kaake, Xiaohui Fang, Mazharul Maishan, Michael A. Matthay, Nevan J. Krogan, Melanie Ott

## Abstract

Severe acute respiratory syndrome coronavirus 2 (SARS-CoV-2) primarily targets the respiratory epithelium, yet severe disease features diffuse lung injury and hyperinflammatory syndromes driven by dysregulated immune activation. Emerging evidence indicates that resident and infiltrating immune cells in the lung can encounter the virus early in infection and, under specific conditions, become infected (*1, 2*). This process amplifies local inflammation and facilitates viral propagation in the lower airways and distal lung regions (*3, 4*), where angiotensin converting enzyme 2 (ACE2) expression is limited (*5, 6*). However, how epithelial and immune cell compartments interact to produce the hallmark pulmonary pathology of SARS-CoV-2 infection remains unresolved. Here we show that secreted ORF8, a SARS-CoV-2 accessory protein, drives inflammatory lung pathology by increasing macrophage permissiveness to infection, triggering pyroptosis, and amplifying viral replication in alveolar epithelial cells. Co-culture of macrophages with human alveolar type II (AT2) cells overrides ORF8’s previously reported inhibition of AT2 infection (*7, 8*), restoring productive viral replication. In vivo, IL-17RA blockade counteracts ORF8 activity, lowering viral burden and attenuating pulmonary inflammation and fibrosis. These findings reveal a paracrine role for ORF8 in reprogramming macrophages, thereby establishing a feedforward proviral circuit that accelerates lung pathology in COVID-19 and are clinically relevant given the recurrent emergence of SARS-CoV-2 variants with either intact or deleted ORF8 since the beginning of the pandemic.

## Main Text

SARS-CoV-2 continues to circulate worldwide despite widespread vaccination and immunity acquired through prior infection (*9, 10*). Within the *Betacoronavirus* genus, the open reading frame 8 (ORF8) is among the most variable genomic regions, frequently undergoing recombination events (*11–14*). ORF8 encodes a secreted accessory protein linked to viral pathogenicity, and serum ORF8 levels in patients with COVID-19 correlate with disease severity (*14–19*). Variants lacking ORF8, such as the Δ382 strain identified in Singapore, are associated with attenuated disease and reduced hypoxia compared to wild-type virus (*20–23*). *In vitro*, ORF8 has been reported to suppress type I interferon signaling, downregulate MHC-I expression, and disrupt host epigenetic control via histone H3 mimicry (*24, 25*). Conversely, ORF8 has also been shown to interfere with viral assembly (*7, 8*), limit new virion production, and bind extracellularly to interleukin-17 receptor A (IL-17RA) (*26*), a mediator of pro-inflammatory signaling. These diverse and sometimes contradictory findings leave the contribution of ORF8 to viral spread and immunopathology mostly unresolved.

Macrophages are central sentinels of the respiratory immune barrier and play a pivotal role in shaping the host response to SARS-CoV-2 (*1*). Severe COVID-19 is characterized by macrophage-driven hyperinflammation (*27–34*), with increased infiltration of macrophages into the lungs (*35, 36*) and genetic associations link inflammasome and pyroptosis pathways to disease severity (*37*). Previous studies have demonstrated that blood monocytes and pulmonary macrophages can be infected by SARS-CoV-2, leading to pyroptotic cell death and increased production of inflammatory cytokines (*30, 38*). However, whether macrophages serve merely as amplifiers of inflammation or as direct targets of productive infection remains a matter of ongoing debate (*3, 30, 39–41*). Here, we tested the hypothesis that ORF8 modified the function of macrophages and their sensitivity to SARS-CoV-2 viral infection.

### Soluble ORF8 protein enhances the uptake of SARS-CoV-2 VLPs by macrophages

To investigate the effect of extracellular ORF8 on SARS-CoV-2 entry into macrophages, we incubated macrophages with virus-like particles (VLPs) (*42*) that incorporated SARS-CoV-2 Spike, Nucleocapsid, E protein, M protein, and the firefly luciferase mRNA (**Fig. 1A**). We found that primary monocyte-derived macrophages (MDMs) take up VLPs irrespective of their polarization into unstimulated macrophages (M0), classically activated macrophages (M1), or alternatively activated macrophages (M2) (**Fig. S1A**). Although VLP-derived luciferase signals in MDMs were substantially lower than those observed in 293T-ACE2/TMPRSS2 cells (**Fig. S2A**), exogenous ORF8 enhanced VLP entry into all MDM subsets in a concentration-dependent manner, with maximal enhancement observed at 1 μg/mL (**Fig. 1B**). Previous studies have reported circulating ORF8 concentrations of approximately 400 ng/mL in patients with severe COVID-19 (*19*); however, the local concentration of ORF8 within infected tissue microenvironments remains unknown. Therefore, we examined a broad concentration range of recombinant ORF8 (0.1–3 μg/mL) in our in vitro assays.

**Fig. 1:**
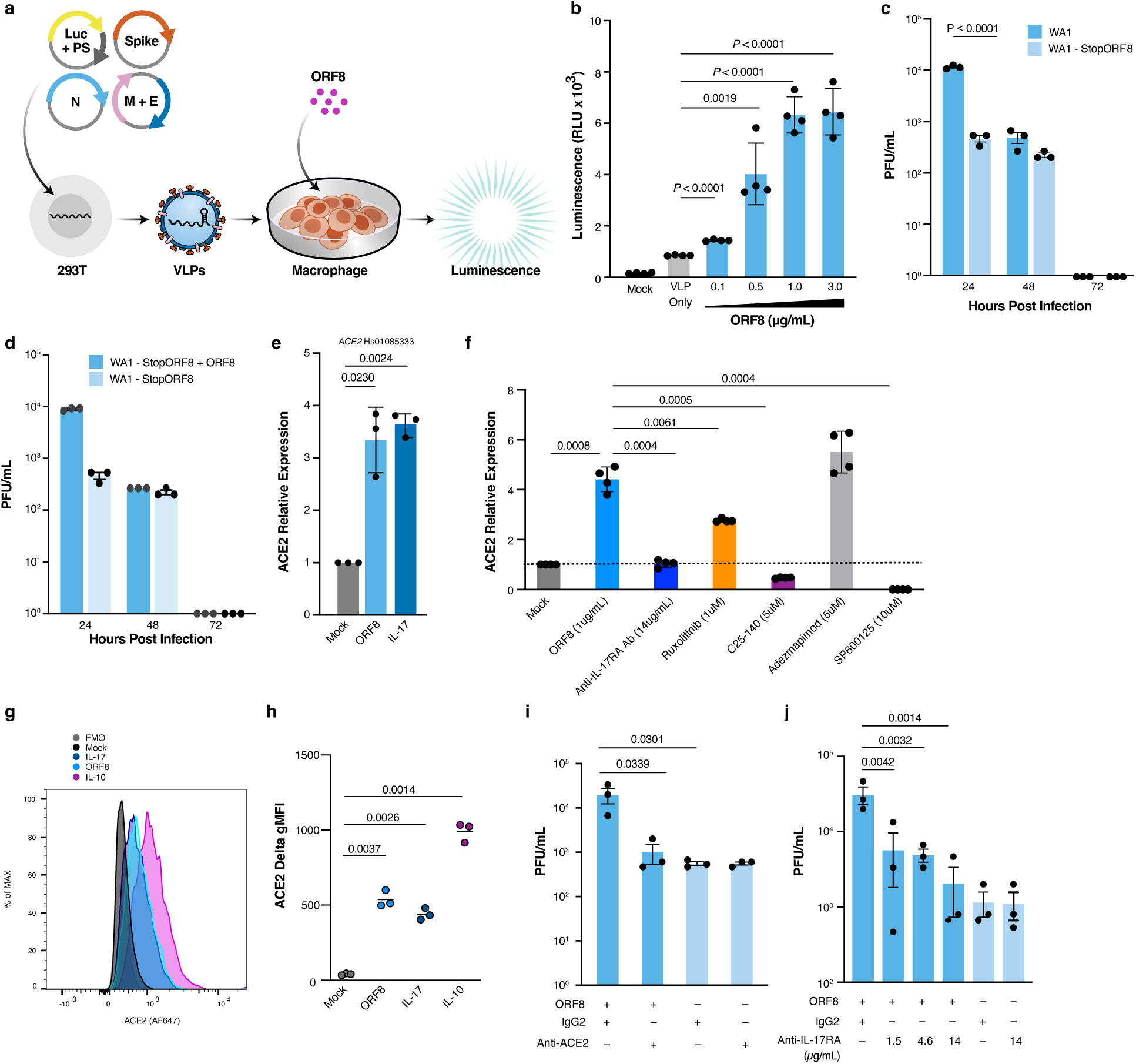
VLP and virion uptake into macrophages is enhanced by soluble ORF8 protein. (A) Schematic of VLP generation and infection of primary monocyte-derived macrophages (MDMs) in the presence of extracellular ORF8. (B) MDMs were treated with increasing concentrations of ORF8 for 48 h and infected with luciferase-expressing SARS-CoV-2 VLPs. Luciferase activity was measured 24 h later. (C and D) MDMs were infected with SARS-CoV-2 WA1 or WA1-StopORF8 (C), or WA1-StopORF8 in the presence or absence of recombinant ORF8 (D). Viral titers in culture supernatants were determined by plaque assay at the indicated time points. (E) Expression of the indicated genes in MDMs treated with ORF8 or IL-17 was quantified by qPCR. (F) MDMs were pretreated with brodalumab (anti–IL-17RA), ruxolitinib, C25-140, adezmapimod, or SP600125 prior to ORF8 stimulation, and ACE2 mRNA expression was measured by qPCR. (G and H) Surface ACE2 expression was quantified by flow cytometry following treatment with ORF8, IL-17, or IL-10. DeltaMFI was calculated by subtracting fluorescence-minus-one (FMO) control values from stained samples. (I and J) MDMs were treated with ORF8 alone or in combination with anti-ACE2 antibody (I) or brodalumab (J), followed by infection with WA1-StopORF8. Viral production was quantified by plaque assay 24 h after infection. Data are presented as mean ± SD. Statistical significance was determined by one-way ANOVA with Kruskal–Wallis and Dunn’s multiple-comparison tests (B, E–J) or two-tailed Welch’s t test (C and D).

### Virally-produced or externally provided ORF8 transiently increases the production of infectious viral particles by infected macrophages

To assess whether infected macrophages can produce new viral particles, we first exposed MDMs to either a SARS-CoV-2 WA1 replicon containing a luciferase reporter gene or to replication-competent SARS-CoV-2 WA1. In the replicon system (*43*). luminescence revealed a significant increase in RNA replication 24 h post-infection. (**Fig. S1C**). In infections with the replication-competent virus, the presence of double-stranded RNA (dsRNA) was detected inside the macrophages by immunostaining as early as 10 h post-infection, and of nucleocapsid protein by 24 h (**Fig. S1D**). We next asked whether virally-produced ORF8 influences viral production. To this end, we infected MDMs with either the replication-competent SARS-CoV-2 WA1 or a mutant strain with a stop codon inserted in the ORF8 gene sequence (StopORF8) at a multiplicity of infection (MOI) of 2 and measured the amount of infectious particles released in the supernatant via a plaque assay. At 24 h post-infection, the wild-type virus expressing ORF8 produced 10 times more viral particles than the StopORF8 virus (**Fig. 1C**). This production excess rapidly declined after 48 h post-infection and by 72 h post-infection, virion production in both WA1- and WA1-StopORF8-infected cells completely ceased (**Fig. 1C**). To test whether externally provided ORF8 could also increase viral production, MDMs were infected with WA1-StopORF8 in the presence of 1 μg/mL full-length extracellular ORF8. Under these conditions, viral particle production at 24 h post-infection increased approximately 10-fold compared with WA1-StopORF8 infection alone, reaching levels comparable to those observed with wild-type WA1 infection (**Fig. 1D**). To evaluate whether ORF8-dependent enhancement of viral production could also be observed in a naturally occurring ORF8-deficient SARS-CoV-2 variant, we repeated these experiments with the XBB.1.5 strain, which carries a G8 nonsense mutation in ORF8 (*44*), in the presence or absence of 1 μg/ml extracellular ORF8. The addition of soluble ORF8 resulted in higher viral particle production up to at least 48 h post-infection, but by 72 h post-infection, viral particle production was undetectable regardless of the presence or absence of extracellular ORF8. (**Fig. S1E**).

### Extracellular ORF8 upregulates ACE2 expression on the macrophages’ cell surface via IL-17RA

To elucidate the mechanism by which ORF8 promotes virion entry into macrophages, we treated MDMs with ORF8 and then measured candidate entry pathway genes by RT-qPCR (**Fig. 1E and S1F**). Extracellular ORF8 treatment induced statistically significant increases in expression of the viral receptor gene *ACE2* (angiotensin-converting enzyme 2) (*45*) (**Fig. 1E**). To identify signaling pathways involved in ORF8-mediated activation of ACE2 transcription in MDMs, cells were stimulated with extracellular ORF8 for 3 h followed by RNA-seq analysis (**Fig. S2, A and B**). Transcriptomic profiling revealed robust induction of inflammatory and chemotactic programs strongly associated with NF-κB and AP-1 signaling. Among the most highly upregulated genes were canonical NF-κB target genes, including IL1B, TNF, CXCL1, CXCL2, CXCL3, CXCL8, CCL3, CCL4, CCL2, TRAF1, PTX3, and TNFAIP6, consistent with activation of a broad proinflammatory transcriptional response. In parallel, multiple negative-feedback regulators of NF-κB signaling, including NFKBIA, NFKBIZ, ZC3H12A, and IER3, were also induced, suggesting engagement of compensatory mechanisms that limit excessive inflammatory activation (**Fig. S2A**). The induction of EDN1, OSM, JAG1, and G0S2 further supported activation of stress-responsive transcriptional programs downstream of NF-κB/AP-1 signaling.

Notably, these transcriptional features were also consistent with activation of IL-17-associated signaling pathways (**Fig. S2B**). Collectively, these findings indicate that ORF8 rapidly activates innate immune programs in human macrophages characterized by coordinated induction of inflammatory cytokines, chemokines, and NF-κB/AP-1-associated regulatory networks. To further define the temporal dynamics of NF-κB and AP-1 activation induced by ORF8, we performed NanoLuc reporter assays in MDMs transduced with NF-κB- or AP-1-responsive reporter lentiviruses (**Fig. S2, C and D**). Cells were stimulated with ORF8 (1 μg/mL), IL-17 (100 ng/mL), ORF8 plus IL-17, or positive controls consisting of LPS (100 ng/mL) for NF-κB reporter assays and PMA (40 nM) for AP-1 reporter assays. Culture supernatants were collected at 5, 10, and 24 h post-stimulation, and extracellular NanoLuc activity was quantified. NF-κB reporter activity in ORF8-stimulated cells was rapidly induced at 5 h but declined to near-baseline levels by 24 h. IL-17 stimulation produced a similar temporal profile. Notably, combined stimulation with ORF8 and IL-17 resulted in strong early NF-κB activation at 5 h, followed by a rapid decline at 10 h and near-complete resolution by 24 h (**Fig. S2C**). In contrast, AP-1 reporter activity exhibited a distinct kinetic profile. Both ORF8 and IL-17 induced relatively modest AP-1 activation at 5 h, which progressively increased over time and reached maximal levels at 24 h (**Fig. S2D**). Co-stimulation with ORF8 and IL-17 similarly resulted in sustained AP-1 activation without the decline observed in NF-κB signaling. Collectively, these findings suggest that ORF8 induces transient NF-κB activation but sustained AP-1 signaling in human macrophages. Building on these findings, we next examined whether ORF8-induced ACE2 expression was functionally linked to IL-17-associated inflammatory signaling pathways (**Fig. 1F**). MDMs were stimulated with recombinant ORF8 (1 μg/mL) in the presence or absence of an anti-IL-17RA antibody (brodalumab), the JAK–STAT inhibitor ruxolitinib, the TRAF6 inhibitor C25-140, the p38 inhibitor adezmapimod, or the JNK inhibitor SP600125, followed by quantification of ACE2 mRNA expression. ORF8-induced ACE2 expression was strongly suppressed by IL-17RA blockade, TRAF6 inhibition, and JNK inhibition, whereas JAK–STAT inhibition produced a more modest reduction (**Fig. 2F**). These findings demonstrate that ORF8 promotes ACE2 transcription primarily through an IL-17RA–TRAF6–JNK signaling axis, consistent with sustained AP-1 activation observed in the reporter assays (**Fig. S2D**). Flow cytometric analysis demonstrated that ORF8-treated MDMs exhibited increased ACE2 surface expression compared with untreated controls (**Fig. 1, G and H**). As a positive control, we also confirmed the previously reported IL-10-mediated upregulation of ACE2 expression (*46*). Since ORF8 interacts with the IL-17RA receptor (*19, 26*), we also incubated MDMs with IL-17. This treatment led to an increase in *ACE2* mRNA expression (**Fig. 1E**) as well as elevated ACE2 protein levels on the cell surface (**Fig. 1, G and H**), mirroring the effects observed with extracellular ORF8.

**Fig. 2:**
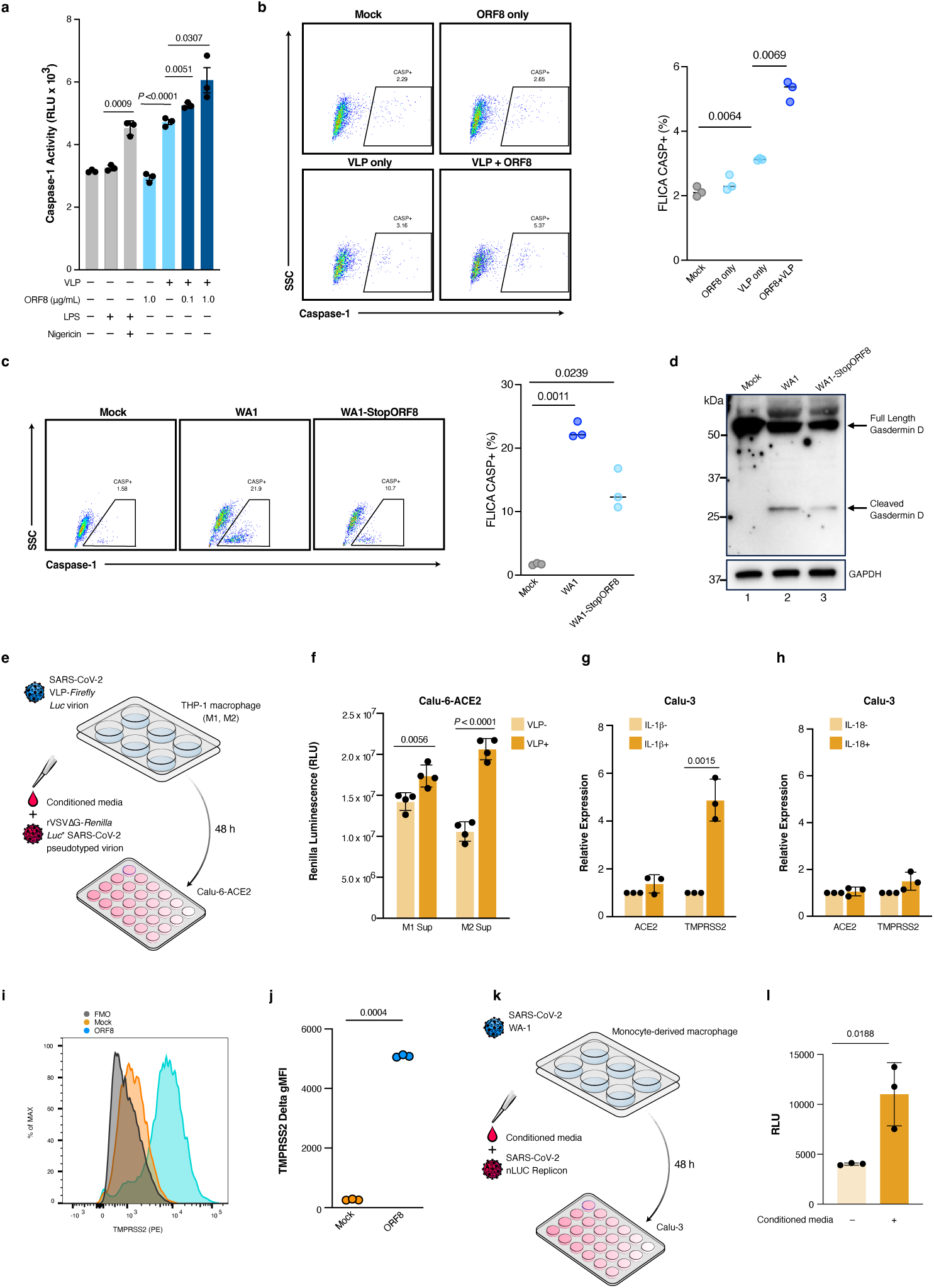
Extracellular ORF8 together with VLP uptake induces pyroptosis in macrophages. (A) MDMs were treated with ORF8, VLPs, or both, and extracellular caspase-1 activity was measured. LPS plus nigericin served as a positive control. (B) Caspase-1 activation in MDMs exposed to VLPs with or without ORF8 was quantified by flow cytometry using FAM-YVAD-FMK staining. Representative plots and summary data are shown. (C) Caspase-1 activation in MDMs infected with SARS-CoV-2 WA1 or WA1-StopORF8. (D) Western blot analysis of adherent and non-adherent cells following infection with WA1 or WA1-StopORF8. (E and F) Experimental design and results of a macrophage-to-epithelial signaling assay. THP-1–derived macrophages were exposed to SARS-CoV-2 VLPs, and conditioned media were transferred to Calu-6-ACE2 cells together with SARS-CoV-2 Spike–pseudotyped reporter virions to assess infectivity. (G and H) Expression of the indicated genes in Calu-3 cells treated with IL-1β or IL-18 was quantified by RT-qPCR. (I and J) TMPRSS2 surface expression in Calu-3 cells following IL-1β treatment was measured by flow cytometry. (K and L) MDMs were infected with replication-competent SARS-CoV-2 WA1, and conditioned media were transferred to Calu-3 cells together with a SARS-CoV-2 NanoLuc replicon to evaluate epithelial cell susceptibility. Data are presented as mean ± SD. Statistical significance was determined by one-way ANOVA with Kruskal–Wallis testing (A) or two-tailed Student’s t test (B, G–L).

To determine whether ORF8-mediated enhancement of ACE2 expression is essential for SARS-CoV-2 infection of macrophages, we assessed de novo viral production of WA1-StopORF8 in the presence or absence of extracellular ORF8, with or without anti-ACE2 antibody AF933, which has previously been reported to inhibit ACE2-dependent viral entry (*47–49*). In the absence of extracellular ORF8, treatment with the anti-ACE2 antibody had no significant effect on viral production. By contrast, in the presence of extracellular ORF8, anti-ACE2 antibody treatment significantly reduced viral production compared with no-antibody controls, decreasing it to levels comparable to those observed in the absence of extracellular ORF8 (**Fig. 1I**). Using the same approach, we next examined whether extracellular ORF8 promotes macrophage infection through the IL-17RA receptor. The addition of brodalumab (anti-IL-17RA antibody) inhibited extracellular ORF8–mediated enhancement of viral particle production in macrophages in a concentration-dependent manner (**Fig. 1J**). These findings indicate that extracellular ORF8 promotes the entry of both VLPs and virions into macrophages by upregulating ACE2 expression on the macrophage cell surface via IL-17RA.

### Extracellular ORF8 combined with VLP uptake induces pyroptosis in macrophages

The drastic decrease in viral production 48 h after infection led us to hypothesize that ORF8-assisted infection induces pyroptosis in macrophages. To test this hypothesis and assess whether ORF8 influences pyroptosis, we evaluated caspase-1 activity (*50*) in the supernatants of MDMs exposed to VLPs with or without soluble ORF8. Whereas ORF8 alone did not activate caspase-1, the combination of VLPs and ORF8 strongly induced caspase-1 activation (**Fig. 2A**).

Furthermore, ORF8 enhanced caspase-1 activity in a dose-dependent manner, with stronger activation observed at 1 μg/mL compared with 0.1 μg/mL ORF8. Increased extracellular caspase-1 activity was used as a surrogate marker of inflammasome-associated pyroptotic cell death, suggesting activation of inflammasome-mediated pyroptotic pathways in macrophages following ORF8-assisted viral uptake. To quantify this effect, we labeled activated caspase-1 using FAM-YVAD-FMK (FLICA), a fluorescent inhibitor probe that covalently binds to active caspse-1, and quantified live caspase-1-activated cells by flow cytometry. The addition of VLPs and ORF8 to MDMs increased the percentage of caspase-1-activated cells by 1.5-fold compared to VLPs alone (**Fig. 2B**). These findings indicate that while extracellular ORF8 by itself does not trigger pyroptosis in macrophages, in combination with VLPs it enhances VLP uptake by macrophage, thereby lowering the threshold for pyroptosis. In a similar experiment using the replication-competent SARS-CoV-2 strains WA1 and WA1-StopORF8, infection with WA1 induced approximately twofold more caspase-1-activated cells than infection with WA1-StopORF8 (**Fig. 2C**). To further assess pyroptotic cell death, both adherent cells and floating cells in the culture supernatant were collected after overnight infection and analyzed by Western blotting. Cleaved gasdermin D was more prominently detected in WA1-infected samples than in WA1-StopORF8-infected samples, further supporting enhanced pyroptotic signaling mediated by ORF8 (**Fig. 2D**).

### IL-1β, a product of macrophage pyroptosis, upregulates TMPRSS2 in lung epithelial cells

In severe SARS-CoV-2 lung lesions, cytokine activation is amplified through crosstalk between epithelial and resident immune cells, with pyroptotic death of bystander myeloid cells proposed to drive cytokine production by epithelial cells (*2, 4*). We aimed to define the mechanistic significance of macrophage pyroptosis, potentiated by ORF8 during SARS-CoV-2 infection, in shaping epithelial cell pathology within the lung. To assess the potential impact of pyroptosis following macrophage infection, we exposed a lung epithelial cell line (Calu-6-ACE2) to the supernatant of infected macrophages before infecting it with SARS-CoV-2 pseudotyped virions. THP-1-derived macrophages were exposed to the VLP described above harboring firefly luciferase mRNA with or without the addition of soluble ORF8, and after 48 h, the macrophage supernatant was mixed with rVSVΔG-Renilla luciferase SARS-CoV-2 pseudotyped virions (*51*) and applied to Calu-6-ACE2 cells. To exclude potential signal contamination arising from carryover of the original firefly luciferase-containing VLPs present in the conditioned media, pseudotyped virions carrying a distinct Renilla luciferase reporter were intentionally used.

Renilla luminescence was quantified 24 h later as a measure of pseudotyped virion entry (**Fig. 2E**). Supernatant from both classically activated macrophages (referred to as the M1phenotype) and alternatively activated macrophages (M2) infected by the SARS-CoV-2 VLP, enhanced virion entry into Calu-6-ACE2 cells (**Fig. 2F**). To test whether early cytokines IL-1β and IL-18 mediate macrophage-driven remodeling of epithelial surface receptor expression, we treated Calu-3 cells with IL-1β or IL-18 (caspase-1-matured). IL-18 had no effect on ACE2 or TMPRSS2 mRNA, while IL-1β significantly upregulated TMPRSS2 transcripts compared with vehicle (**Fig. 2, G and H**). Furthermore, flow cytometric analysis of IL-1β-stimulated Calu-3 cells demonstrated increased TMPRSS2 surface expression (**Fig. 2I**). Calu-3 were used after culture on Transwell inserts under air-liquid interface conditions to induce epithelial polarization (*52*). To confirm the infection-enhancing effect of supernatants derived from infected macrophages on lung epithelial cells, MDMs were exposed to replication-competent SARS-CoV-2 WA1, followed by extensive washing to remove residual input virus. Culture supernatants were collected 48 h post-infection and added to Calu-3 cells together with a SARS-CoV-2 replicon carrying a NanoLuc reporter. NanoLuc activity measured 24 h later revealed significantly higher relative luminescence units in cells treated with conditioned media from infected macrophages, indicating enhanced viral replication in epithelial cells (**Fig. 2, K and L**).

### Co-culture with macrophages allows the replication of ORF8-expressing SARS-CoV-2 in AT2 lung epithelial cells

To investigate epithelial cells-macrophage interaction during SARS-CoV-2 infection and tease apart the role of ORF8 under more physiological conditions, we infected mono-and co-cultures of MDMs and primary lung epithelial cells with WA1 or WA1-stopORF8 viruses. Initially, MDMs were spread at the bottom of the Transwell plates, and the virus was added at an MOI of 1. Viral mRNA was consistently higher in WA1-infected cells than in WA1-stopORF8-infected cells, supporting a proviral role of ORF8 in macrophage infection (**Fig. 3A**). Plaque assays further demonstrated that infectious viral particles were detectable only in the supernatants of WA1-infected cultures at 24 h post-infection, whereas infectious virus was undetectable in WA1-StopORF8-infected cultures at 24, 48, and 72 h post-infection (**Fig. 3A**). In contrast, in monocultures of primary alveolar type 2 (AT2) cells (*53, 54*), infection with WA1-stopORF8 resulted in higher levels of viral RNA than infection with WA1, confirming previous reports of an antiviral role of ORF8 in epithelial cell infection (*7, 55*) (**Fig. 3B**). Consistent with these findings, plaque assays demonstrated that WA1-StopORF8 infection also produced higher titers of infectious viral particles than WA1 infection in AT2 monocultures (**Fig. 3B**). Finally, we infected co-cultures in which MDMs were grown on the bottom and AT2 cells on the upper chamber of the Transwell plate. In co-culture conditions with macrophages (**Fig. 3C**), AT2 cells exhibited a 20-(*p* = 0.0468) and 7-fold (*p* = 0.0246) increase in viral mRNA expression at 48-and 72-h post-infection, respectively, compared to AT2 cells cultured alone (**Fig. 3B**). Consistent with these findings, plaque assays demonstrated that WA1 infection under co-culture conditions produced the highest titers of infectious viral particles among all experimental conditions tested (**Fig. 3C**). These results suggest that ORF8’s promotion of macrophage infection in allowing efficient viral replication in lung epithelial cells.

**Fig. 3:**
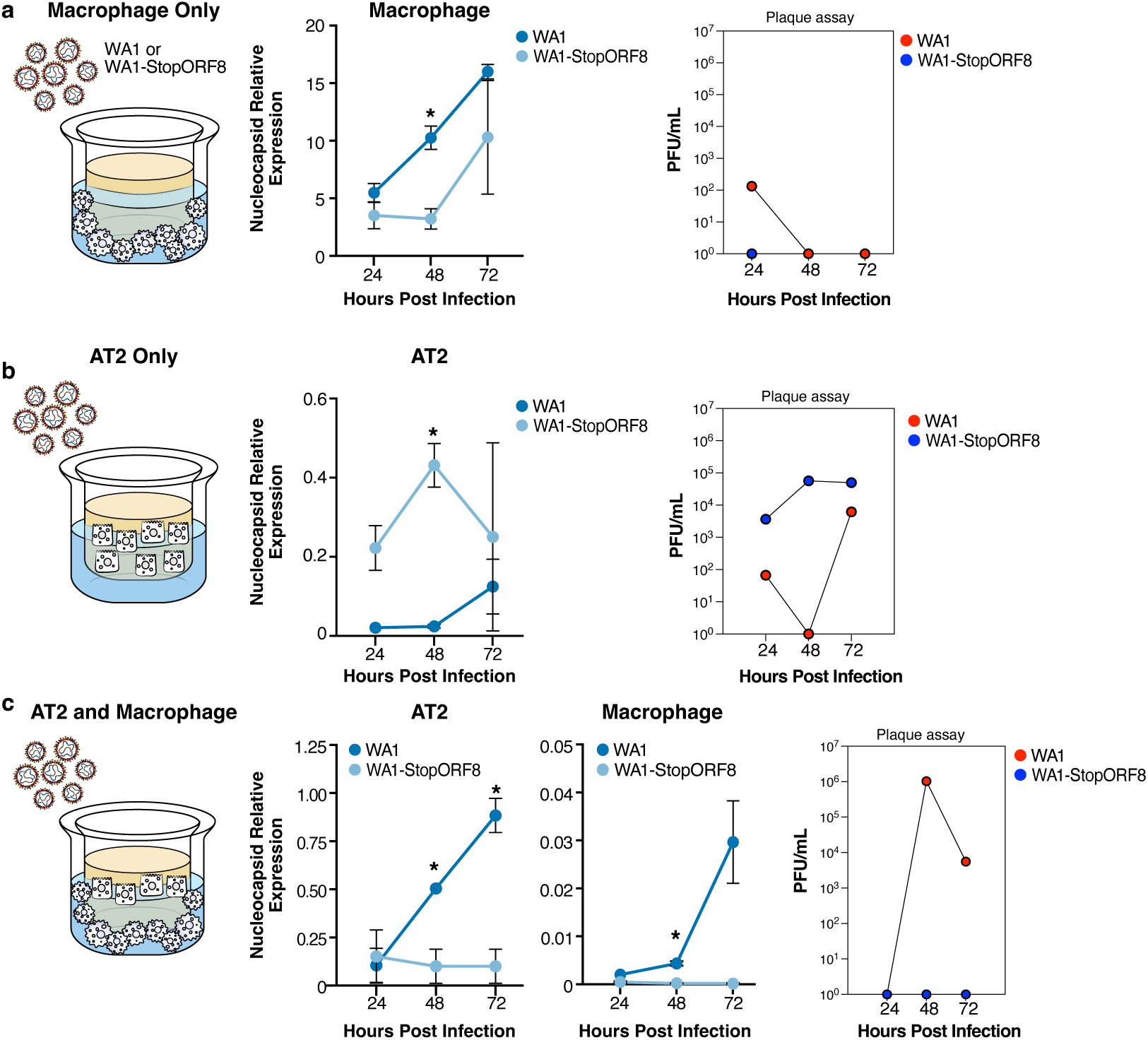
Viruses containing ORF8 enable SARS-CoV-2 to fully replicate in AT2 cells and macrophages. (A) MDMs were seeded on the bottom of 24-well Transwell plates and replication-competent SARS-CoV-2 WA1 or WA1-StopORF8 was added at MOI = 1. After 2 h, the cells were washed and flesh medium added. After 24, 48, and 72 h, the cells were collected and lysed. SARS-CoV-2 *N* mRNA in cell lysate was quantified by RT-qPCR normalized to HPRT. Plaque assays were performed using culture supernatants collected from each well of the WA-1 and WA-1-StopORF8 groups. Data are presented as mean values. (B) Primary human alveolar type 2 (AT2) cells were seeded in the upper chamber of 24-well Transwell plates. The two virus strains were added and analyzed as in (A). (C) MDMs and AT2 cells were co-cultured, with MDMs in the bottom and AT2 cells in the upper chamber of a 24-well Transwell plate. The two virus strains were added, washed after 2 h, and cells were analyzed separately by qPCR at 24, 48, and 72 h later. Data are presented as 2^-ΔCt^ (means ± SD of three independent experiments). Significance was assessed by two-tailed Student’s t-test. \**p* < 0.05.

### Extracellular ORF8 shifts macrophages to an M2-like phenotype

IL-17 exerts dual functions, activating macrophages to protect the homeostatic state of tissues while also promoting inflammatory disease (*56, 57*). We hypothesized that ORF8 reprograms macrophage phenotypes through mechanisms extending beyond modulation of ACE2 expression. To assess whether ORF8 affects macrophage phenotypes, we exposed unstimulated macrophages (referred to as M0), classically activated macrophages (M1), and alternatively activated macrophages (M2) to extracellular ORF8 and performed proteomic profiling at 12 and 48 h, quantifying 5723 total proteins in at least one condition (**Fig. 4A, Table S**1). M0 macrophages showed increased CD206, a representative marker of M2 macrophages, at 48 h after ORF8 exposure and reduced DDX60 and IRF3, components of the RIG-I–dependent type-I interferon pathway (**Fig. 4B, left panel**). In M1 macrophages, CCR7, a representative marker of M1 macrophages, was downregulated, whereas HMOX1, MMP9, CHI3L1, and FGL2, proteins enriched in M2 macrophages, were upregulated (**Fig. 4B, middle panel**). In M2 macrophages, ORF8 preferentially amplified an M2-leaning profile, increasing mitochondrial/OXPHOS proteins (MRPL23, TIMM8A, COX5B, CYCS), antioxidant/stress-response factors (PRDX1, HPX, AK4), and immunoregulatory components (CSF1, LGALS3, CASP9, UBE2N), together with lipid/metabolic–stress proteins (APOC3, SRI), without broad induction of inflammatory markers (**Fig. 4B, right panel**). Across activation states, nine proteins—MMP9, APOC3, ITIH4, PVR, CASP9, CSF1, CYP1B1, MRPL23, and RUFY2—were upregulated in at least two macrophage states, mapping to pathways of tissue repair/remodeling, lipid/xenobiotic metabolism, mitochondrial function, immune regulation, cell-death signaling, and vesicular trafficking (**Fig. 4, B and C**). Collectively, these data indicate a shift toward an M2-like profile enriched for tissue-repair and lipid-metabolism pathways, with inflammatory signatures attenuated. Gene Ontology (GO) analysis indicated an enrichment in multiple immune-response pathways for M1 macrophages at 48 h post extracellular ORF8 exposure (**Fig. S3, Table S2**), therefore we analyzed polarity-associated mRNA changes in M1 cells using reverse transcription quantitative PCR (RT-qPCR). Exposure to extracellular ORF8 increased mRNA levels of M2-associated genes—*ARG1*, *VEGFA*, *IL10*, and *MRC1* (CD206)—whereas transcripts of interferon-pathway genes *IRF7*, *OAS1*, and *CXCL10* were reduced (**Fig. 4D**). For functional assays, phagocytosis was evaluated using pHrodo™ *E. coli* bioparticles. M1 macrophages pre-exposed to ORF8 exhibited increased fluorescence relative to controls, consistent with greater phagosome acidification/uptake (**Fig. 4E**). Furthermore, with lipopolysaccharide (LPS) at 1 ng/mL, IL-10 secretion by ORF8-exposed M1 macrophages did not differ significantly from unexposed controls; at 10 and 100 ng/mL it increased 9.2-fold (*p* = 0.0005) and 3.6-fold (*p* = 0.0026), respectively, as quantified using an enzyme-linked immunosorbent assay (ELISA) of culture supernatants (**Fig. 4F**).

**Fig. 4:**
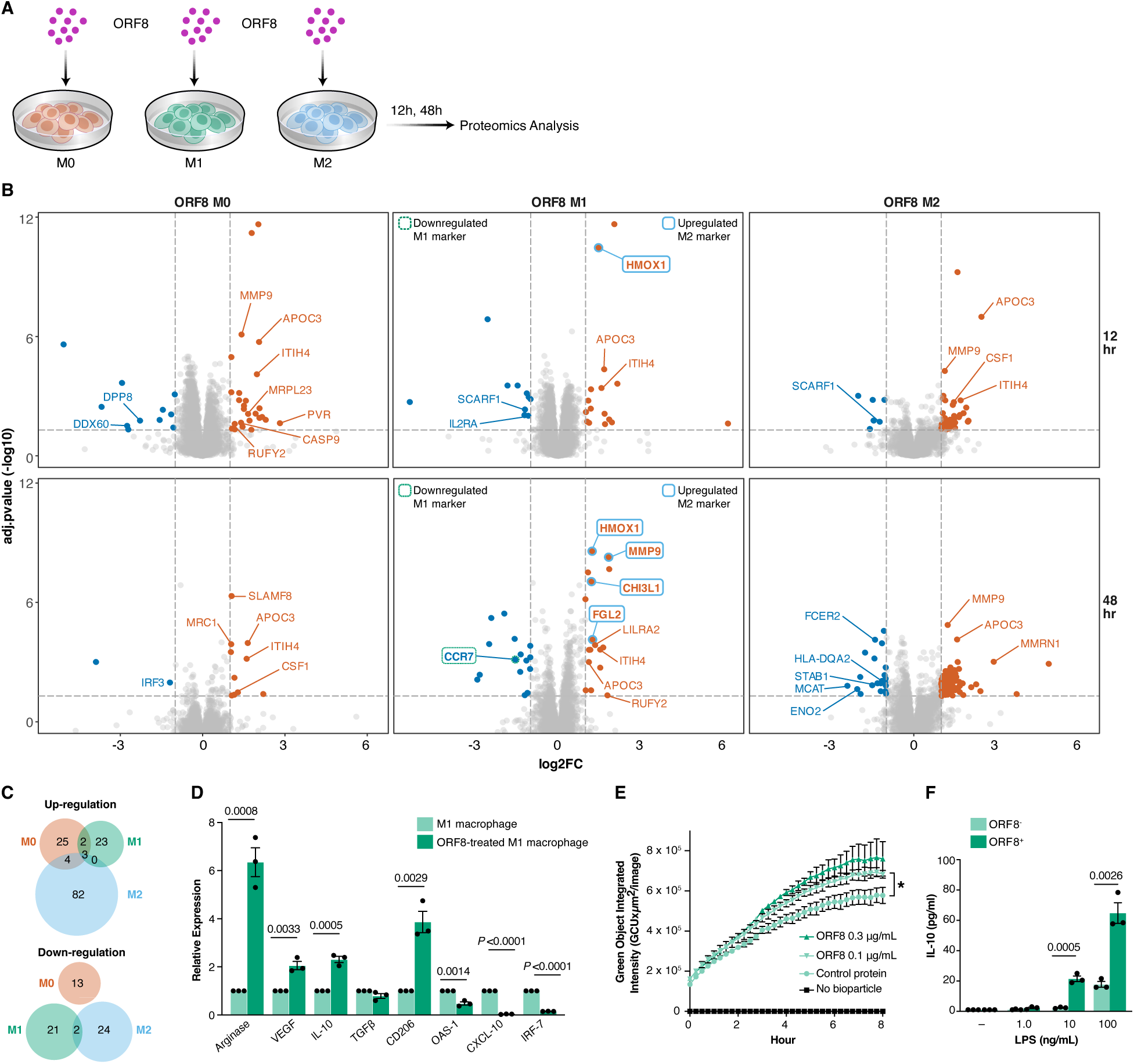
Extracellular ORF8 shifts macrophages to an M2-like phenotype. (A) Schematic diagram of the proteomic analysis performed on MDMs (M0, M1, M2) cultured in the presence of extracellular ORF8. (B)Volcano plots depicting differential protein expression in macrophages of three polarization states (M0, M1, M2) at 12-and 48-h following exposure to ORF8. The x-axis represents log₂ fold change (log₂FC), and the y-axis represents adjusted p-values. Proteins to the right (red) are significantly upregulated, whereas those to the left (blue) are significantly downregulated in response to ORF8. Significance thresholds were defined as adjusted *p* < 0.05 and |log₂FC| > 1. (C) Venn diagrams showing proteins significantly upregulated (upper) or downregulated (lower) in response to ORF8 exposure across three macrophage subtypes. (D) MDMs (M1) were incubated with ORF8 (1μg/mL) or vehicle for 48 h. Expression of the indicated genes was quantified by RT-qPCR normalized to HPRT. (E) MDMs (M1) were cultured at 37 °C for 8 h in the presence of extracellular ORF8 or SARS-CoV-2 N protein (control), together with pHrodo™ *E. coli* bioparticles. Fluorescence was monitored and quantified using the Incucyte live-cell analysis system. \**p* < 0.05. (F) MDMs (M1) was incubated with ORF8 or vehicle for 48 h, then LPS (1, 10, or 100 ng/mL) was added to the cell culture medium. The cell supernatant was harvested 24 h later and tested with IL-10 ELISA assay. Data are presented as fold change relative to unstimulated ± SD of three independent experiments. Significance was assessed by two-tailed Student’s t-test.

### IL17RA antibody limits viral replication and lung pathology in ORF8-SARS-CoV-2-infection mice

To determine whether ORF8 influences viral replication and ACE2 expression in lung cells under *in vivo* conditions, C57BL/6 wild-type mice were infected with mouse-adapted SARS-CoV-2 WA1 (ORF8+), the StopORF8 WA strain, or the StopORF8 together with intraperitoneally administered recombinant ORF8 (ORF8 IP). Two days after infection, lungs were harvested, and macrophages and epithelial cells were isolated by antibody-based selection (**Fig. 5, A and B**). mRNA levels of the viral *N* gene and *Ace2* were quantified by RT-qPCR. In macrophages, *N* gene expression was increased 5.0-fold in WA1-infected mice (*p* = 0.0428) and 5.2-fold in ORF8 IP mice (*p* = 0.0339) compared with StopORF8-infected mice (**Fig. 5B**). *Ace2* expression was also elevated 4.3-fold in ORF8 IP mice (*p* = 0.0327), and WA1-infected mice showed a similar upward trend (**Fig. 5C**). In epithelial cells, *N* gene expression was 10-fold higher in WA1-infected mice (*p* = 0.0038) and 5.6-fold higher in ORF8 IP mice (*p* = 0.0474) compared with StopORF8-infected mice (**Fig. 5E**), whereas *Ace2* expression showed no significant differences between groups (**Fig. 5F**). To evaluate the therapeutic potential of IL-17RA blockade, we repeated these experiments on an additional cohort in which IL17RA antibody or a control IgG was administered intraperitoneally (**Fig. 5G**). In this cohort, infectious viral titers were 5.1-fold higher in the WA1 group (*p* = 0.0008) and 5.8-fold higher in the ORF8 IP group (*p* < 0.001) compared with the StopORF8 group in the absence of brodulamab. IL17RA antibody treatment reduced viral titers by 3.4-fold in the WA1 group (*p* = 0.0035), eliminating the difference from the StopORF8 group, whereas the StopORF8 group itself showed no response to IL17RA antibody. In contrast, the ORF8 IP group exhibited a 7.3-fold reduction in viral titer upon treatment with IL17RA antibody (*p* < 0.0001) (**Fig. 5H**). Histological analysis of lung tissue by haematoxylin–eosin staining and caspase-1 immunohistochemistry (IHC) revealed reduced air space and increased numbers of caspase-1–positive cells, including those in the airways, in the WA1 and ORF8 IP groups compated to the StopORF8 group (**Fig. 5I**), but these differences were abolished after intraperitoneal IL17RA antibody treatment (**Fig. 5J**). The area of caspase-1–positive regions was also elevated in WA1 and ORF8 IP groups and decreased following IL17RA antibody (**Fig. 5K**). In addition, Trichrome staining of WA1-infected lungs showed reduced fibrotic areas after IL17RA antibody treatment on five days after infection (**Fig. 5, L and M**).

**Fig. 5:**
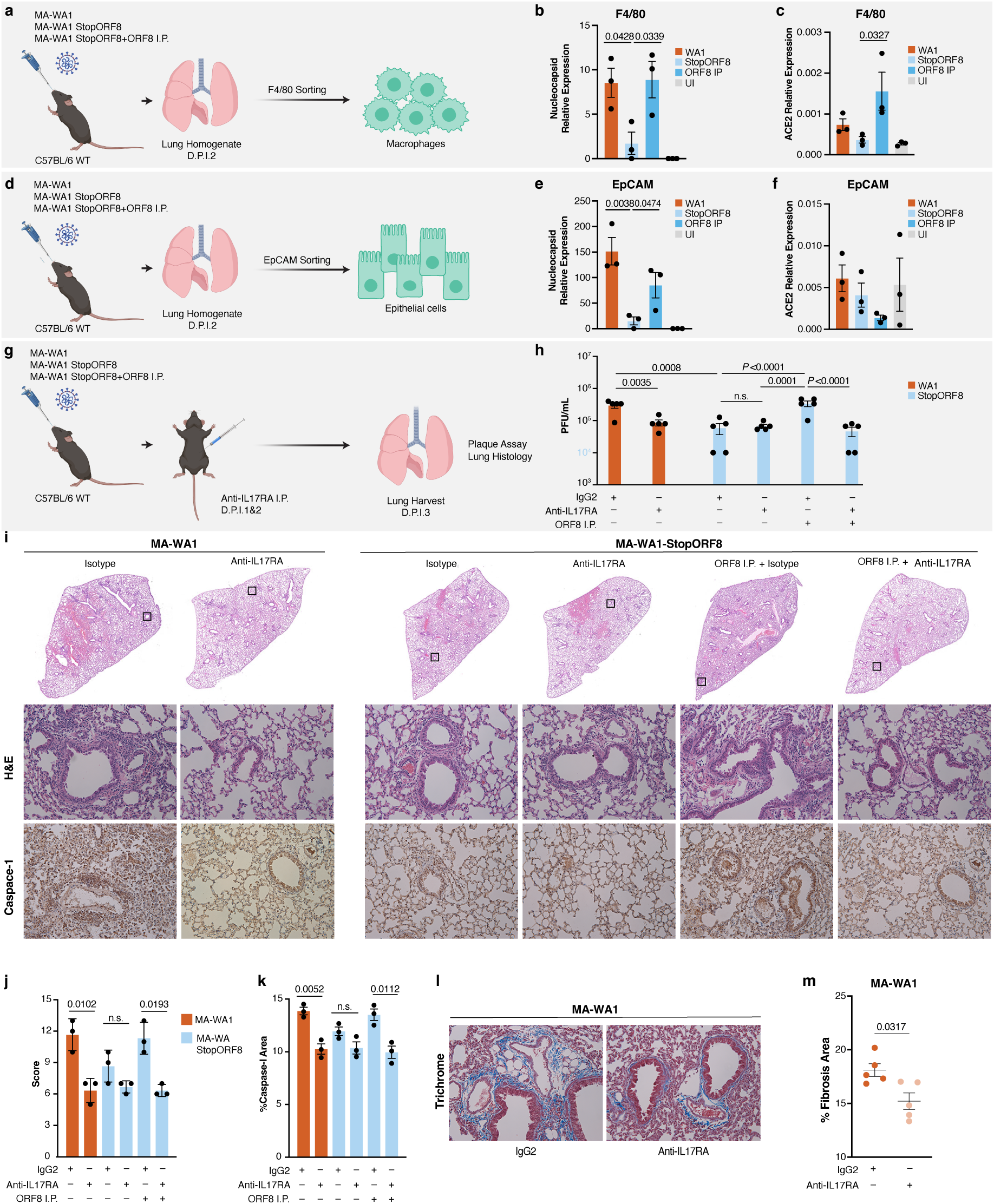
IL17RA Antibody Limits Viral Replication and Lung Pathology in ORF8-positive SARS-CoV-2 Infection. (A-F) C57BL/6 wild-type mice were infected with mouse-adapted WA1 (10^4^ PFU), mouse-adapted (MA) WA1-StopORF8, or WA1-StopORF8 supplemented with 10 μg of extracellular ORF8 administered intraperitoneally (IP) at 1-day post-infection (DPI 1). At 2 DPI, F4/80⁺ macrophages (A) and EpCAM⁺ epithelial cells (D) were isolated by positive selection, and the expression of SARS-CoV-2 *N* (B, E) and *Ace2* (C, F) mRNA was quantified by RT–qPCR. **g**,**h**, C57BL/6J mice were infected with mouse-adapted WA1 (10⁴ PFU), WA1-StopORF8, or WA1-StopORF8 supplemented with IP injection of 10 μg extracellular ORF8 at 1 and 2 DPI. Each group was further divided into two subgroups, receiving either IL17RA antibody (100 μg, IP at DPI 1 and 2) or IgG1 as an isotype control. In the ORF8-treated group, IL17RA antibody was administered IP 2 h after ORF8 injection. At 3 DPI, the lung tissue was homogenized, and infectious viral particles were quantified by plaque assay. Significance was assessed by one-way ANOVA with a Kruskal–Wallis test with Dunn’s multiple comparison tests. (I, J, K) Lungs collected using the same procedure were fixed, stained with hematoxylin and eosin (H&E) and caspase-1 immunohistochemistry (IHC), and caspase-1–positive areas were quantified by image analysis. (L, M) In MA-WA1–infected mice, the extent of fibrosis was assessed by Masson’s trichrome staining at 5 DPI. Significance was assessed by two-tailed Student’s t-test. \**p* < 0.05.

In an independent infection experiment using aged BALB/c mice, which are susceptible to mouse-adapted SARS-CoV-2 infection (*58*), mice infected with MA-WA1-StopORF8 exhibited only mild weight loss. In contrast, MA-WA1-infected mice developed more severe disease, with of 5 animals exceeding the predefined euthanasia threshold of 15% body weight loss (**Fig. S4A**). Consistent with these clinical findings, lung plaque assays revealed significantly higher viral titers in MA-WA1-infected mice compared with MA-WA1-StopORF8-infected mice (**Fig. S4B**). Based on these results, we propose a working model in which ORF8 engages IL-17RA signaling in macrophages to promote viral replication and lung pathology (**Fig. 6**).

**Fig. 6:**
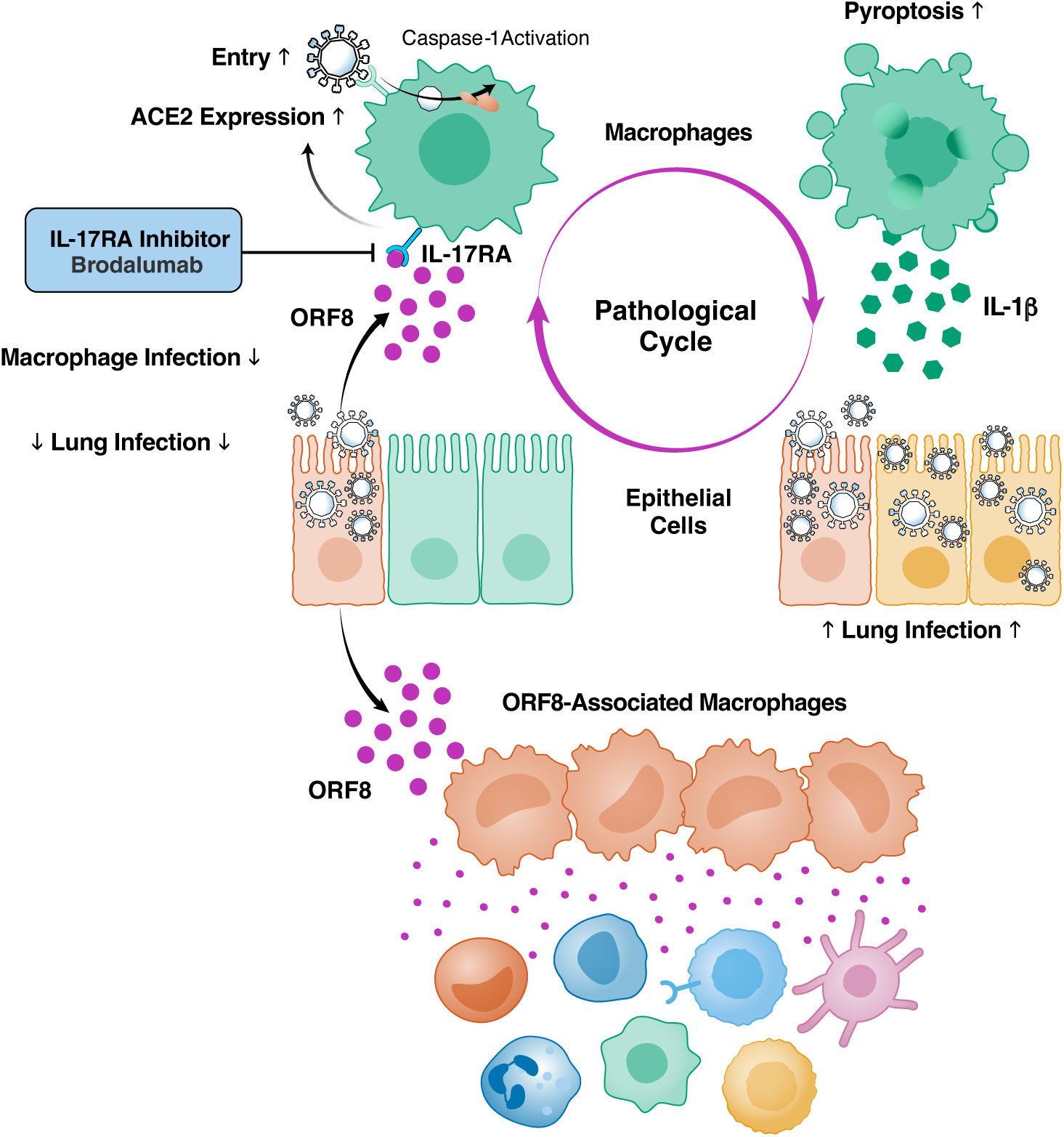
Model of ORF8-driven macrophage reprogramming enhancing SARS-CoV-2 epithelial infection. Secreted ORF8 engages IL-17RA on macrophages, upregulating ACE2 and enhancing viral uptake. Infection triggers caspase-1 activation and pyroptosis, releasing IL-1β that increases epithelial susceptibility to SARS-CoV-2. ORF8 also reprograms macrophages toward an immunosuppressive phenotype. Together, these events create a vicious cycle that amplifies viral replication and disease severity. IL-17RA blockade disrupts this loop and mitigates pneumonia.

## Discussion

Our study identifies macrophages as key cellular targets of SARS-CoV-2 ORF8 and uncovers mechanisms by which ORF8 promotes viral spread while undermining host defense. We show that secreted ORF8 upregulates ACE2 in macrophages, markedly increasing their susceptibility to infection and pyroptotic cell death. This creates a feedforward loop in which infected macrophages enhance infection of epithelial cells, amplifying viral dissemination within the respiratory tract. Mechanistically, our data indicate that ORF8 induces ACE2 expression through an IL-17RA–TRAF6–JNK signaling axis associated with sustained AP-1 activation.

Transcriptomic analyses revealed rapid induction of inflammatory and IL-17-associated gene programs following ORF8 stimulation, while pharmacological inhibition of IL-17RA, TRAF6, or JNK strongly suppressed ORF8-induced ACE2 expression. Notably, our reporter assays demonstrated that ORF8-induced NF-κB activation was rapid but transient, declining to near-baseline levels within 24 h, whereas AP-1 activation progressively increased over time. The rapid attenuation of NF-κB signaling, together with induction of multiple negative-feedback regulators suggests that ORF8-triggered NF-κB activation is tightly self-limited following the initial inflammatory response. In contrast, sustained AP-1 signaling may play a dominant role in maintaining ACE2 transcriptional upregulation. Previous studies have reported that macrophages express minimal or undetectable levels of IL-17RC (*59*), raising the possibility that ORF8 may engage IL-17RA signaling through a noncanonical mechanism. In addition, previous reports have suggested that only non-glycosylated ORF8 is capable of binding IL-17RA (*60*). Because recombinant ORF8 produced in mammalian expression systems likely contains heterogeneous glycosylation states, differences in ORF8 production methods may substantially influence downstream macrophage responses. Indeed, some groups have reported minimal transcriptional responses following ORF8 stimulation of macrophages (*61*). Such discrepancies may reflect differences in ORF8 synthesis and purification methods, macrophage differentiation and culture conditions, or the timing of sample collection following stimulation. How ORF8 functionally interacts with the IL-17 receptor complex in macrophages remains an important subject for future investigation. Consistent with this model, our in vivo experiments demonstrated that blockade of IL-17RA significantly reduced viral replication to levels comparable to those observed with StopORF8 infection and improved disease outcomes.

At the same time, ORF8 reprograms macrophages from an inflammatory M1 profile toward an immunosuppressive M2-like state. Macrophages characterized by high CD206 expression, a hallmark of the M2-like state, are thought to play a role in SARS-CoV-2 pathogenesis (*1, 3, 34*). This shift, reinforced by IL-10, which is known to upregulate ACE2 on macrophages (*46*), mirrors the behavior of tumor-associated macrophages in suppressing local immune responses. Although pyroptotic inflammation and M2-like polarization appear contradictory, together they establish an immunopathological environment that favors viral replication while blunting antiviral immunity. Importantly, pyroptosis is accompanied by dysregulated release of IL-1β, which is known to drive aberrant AT2 self-renewal and promote fibrotic remodeling (*4, 62*). This association raises the possibility that ORF8 contributes to post-acute sequelae of COVID-19 (PASC) by allowing macrophages to sustain inflammation (*63*) beyond the acute phase.

Finally, our co-culture experiments highlight the importance of epithelial–macrophage crosstalk in disease progression. ORF8 not only enhances pyroptosis and viral uptake in macrophages but also increases epithelial permissiveness through macrophage-derived signals. In agreement with our findings, previous work demonstrated synergistic cytokine induction in macrophage–lung epithelial cell co-culture systems in the presence of ORF8 (*60*). These observations are consistent with our proposed pathological cycle in which ORF8-mediated ACE2 upregulation enhances macrophage infection, leading to excessive inflammatory cytokine production that further promotes epithelial cell susceptibility and viral dissemination. Interestingly, the marked discrepancy between intracellular nucleocapsid mRNA levels and the amount of infectious virus detected in macrophage monocultures (Fig. 3A) suggests that only a limited subset of macrophages that take up viral particles supports productive viral replication. In this context, ORF8-mediated enhancement of viral uptake through ACE2 upregulation may critically increase the likelihood of establishing replication-competent infection in macrophages. Our findings may help reconcile previous conflicting reports regarding macrophage infection by SARS-CoV-2.

Together, these findings identify ORF8 as a central driver of epithelial–immune cell interactions in COVID-19, positioning it as both a mechanistic link between viral replication and immune evasion, and a promising therapeutic target.

Several limitations of this study should also be acknowledged. Although macrophage infection was confirmed by immunostaining analyses (Fig. S1D), the overall frequency of infected macrophages remained relatively low and varied substantially among donors. Such variability may reflect differences in donor-specific innate immune states or prior immunological exposures. In particular, the potential contribution of trained innate immune memory to macrophage susceptibility and inflammatory responses following SARS-CoV-2 exposure warrants further investigation. Because ORF8-enhanced infection rapidly induced pyroptotic cell death, accurate quantification of infected macrophages over time was technically challenging.

The rapid loss of infected cells may therefore lead to underestimation of the true extent of macrophage infection and viral replication dynamics.

## Acknowledgments

We are deeply grateful to Dr. Jennifer A. Doudna and Dr. Abdullah M. Syed for graciously sharing the VLP constructs. We thank Donor Network West for the procurement of donor lungs for research, and the donors and their families.

## Funding

National Institutes of Health grant U19AI135990 (MO, NJK)

National Institutes of Health grant U54HL147127 (MAM)

James B. Pendleton Charitable Trust (MO)

Roddenberry Foundation (MO)

Pam and Ed Taft (MO)

Chan Zuckerberg Biohub – San Francisco (MO)

Fast Grants (MO)

Innovative Genomics Institute (MO)

California HIV/AIDS Research Program (YM)

NIH Division of Intramural Research (RKS)

## Author contributions

Conceptualization: YM, RKS, MO

Methodology: YM, RKS, MMontano, TYT, MMK, LS, XF, MMaishan, JT, YZ, RMK, MAM, NJK, MO

Investigation: YM, RKS, MMontano, TYT, JT, YZ, RMK, NJK, MO Visualization: YM, RKS, TYT, YZ, RMK

Funding acquisition: MO, NJK, MAM, YM Project administration: KK

Supervison: MAM, NJK, MO Writing – original draft: YM, MO

Writing – review & editing: YM, MO, KK, RKS, MMontano, TYT, MMK, LS, XF, MMaishan, JT, YZ, RMK, MAM, NJK

## Competing interests

The N.J.K. laboratory has received research support from Vir Biotechnology, F. Hoffmann-La Roche, and Rezo Therapeutics. N.J.K. has a financially compensated consulting agreement with Maze Therapeutics. N.J.K. is the Interim CEO and is on the Board of Directors of Mreza Therapeutics, and he is a shareholder in Mreza Therapeutics, Rezo Therapeutics, Tenaya Therapeutics, Maze Therapeutics, and GEn1E Lifesciences.

## Data, code, and materials availability

All data and code needed to evaluate and reproduce the results in the paper are present in the paper and/or the Supplementary Materials. Engineered cell lines generated in this study are available from the corresponding author upon reasonable request, subject to scientific review and completion of a Material Transfer Agreement (MTA) with Gladstone Institutes. All other data supporting the findings of this study are available within the paper and its Supplementary Materials.

## Supplementary Materials

Materials and Methods

Figs. S1 to S5

Tables S1 and S7

