## Supplemental materials for "Secreted ORF8 reprograms macrophages to enhance SARS-CoV-2 infection of lung epithelial cells"

**The PDF file includes:**

Materials and Methods
Figs. S1 to S5

**Other Supplementary Materials for this manuscript include the following:**

Tables S1 to S7

### Materials and Methods

#### Ethics statement

All research conducted in this study complies with all relevant ethical regulations. All experiments conducted with replication-competent viruses were performed in a certified biosafety level 3 (BSL3) laboratory and experiments were approved by the Institutional

Biosafety Committee of the University of California, San Francisco and the Gladstone Institutes. All protocols concerning animal use were approved (AN203103-00F) by the Institutional Animal Care and Use Committees of the University of California, San Francisco and the Gladstone Institutes and conducted in strict accordance with the National Institutes of Health Guide for the Care and Use of Laboratory Animals. All human samples utilized for this project were de-identified, not used to conduct human subject research, and were therefore IRB exempt.

**ORF8 production**

The ORF8-Flag expression construct was generated by ChemPartner. Recombinant C-terminal Flag-tagged ORF8 protein was expressed in HEK293 cells in a 2-L suspension culture for 7 days and purified using anti-Flag affinity purification followed by Superdex 200 size-exclusion chromatography. The purified protein was formulated in 50 mM Tris, 200 mM NaCl, and 1 mM DTT (pH 8.0). Protein purity was estimated to be approximately 90% by SDS–PAGE analysis. The amino acid sequence of the recombinant ORF8-Flag protein was as follows: MKFLVFLGIITTVAAFHQECSLQSCTQHQPYVVDDPCPIHFYSKWYIRVGARKSAPLIEL CVDEAGSKSPIQYIDIGNYTVSCSPFTINCQEPKLGSLVVRCSFYEDFLEYHDVRVVLDFI AIAGAAADYKDHDGDYKDHDIDYKDDDDK.

**SARS-CoV-2 propagation and infection**

The SARS-CoV-2 isolate USA-WA1/2020 was produced by cloning into a molecular clone expression plasmid, followed by transfection into BHK-21 cells (ATCC) (*43*). The WA1-stopORF8 clone was generated by site-directed mutagenesis as described previously8. These viruses were then cultured in Vero E6 cells (ATCC) capable of expressing human ACE2 and TMPRSS2 receptors on the cell membrane. The cells were supplemented with a medium containing 2% FBS, D-glucose (4.5 g/L), 4 mM L-glutamine, 10 mM nonessential amino acids, 1 mM sodium pyruvate, and 10 mM HEPES. This process involved creating two successive virus stocks.

Three days post-infection, the supernatant, which contained the propagated virus, was filtered through an Amicon Ultra 15 (100-kDa) centrifugal filter (Millipore Sigma) at approximately 4,000 rpm for 20 minutes. The flow-through was discarded, and the virus was resuspended in DMEM. The infectivity titer of SARS-CoV-2 was determined using a plaque assay on Vero E6-ACE2-TMPRSS2 cells in minimal essential medium enhanced with 2% FBS, 4 mM L-glutamine, 0.2% bovine serum albumin, 10 mM HEPES, 0.12% $NaHCO_3$, and 0.7% agar. The multiplicity of infection (MOI) values used in the experiments were based on titers obtained from these plaque assays. All procedures involving live SARS-CoV-2 were conducted in the BSL-3 facility at Gladstone Institutes. This facility is certified by both the CDC and USDA and conforms to the institutional biosafety requirements. For ACE2-targeted infection-inhibition assays, cells were preincubated with an anti-human ACE2 antibody (R&D Systems) or an-isotype-matched human IgG2 control (Caltag Laboratories). For macrophage assays targeting IL-17RA, cells were preincubated with brodalumab (Selleckchem) or an isotype-matched human IgG2 control (Caltag Laboratories).

**Plaque assay**

Supernatants from cell cultures were assessed for the formation of viral particles in both cell and in organoid experiments. In summary, Vero-TMPRSS2 cells were seeded and allowed to incubate overnight. Subsequently, cells were exposed to $10^{-1}$ to $10^{-6}$ dilutions of the respective homogenates or supernatants in serum-free DMEM. Following a 1-hour absorption period, a 2.5% Avicel (RC-591, Dupont) overlay was applied to the wells. After 72 h, the overlay was removed, and the cells were fixed in 10% formalin for 1 hour, followed by staining with crystal violet for the visualization of plaque-forming units (PFU). Data analysis was carried out using GraphPad Prism version 10.2.0.

Production of viral-like particles and assessment of transduction efficiency by luciferase assay

Plasmids CoV2-N (10µg), CoV2-M-IRES-E (5µg), CoV-2 Spike (24ng), and Luc-T20 (15µg) were mixed in a total volume of 2 mL of Opti-MEM (Gibco) (*42*). To this DNA solution, 90 µg of PEI was added, resulting in a final volume of 2 mL in Opti-MEM. The transfection mixture was then allowed to incubate at room temperature for 20 minutes. Following this incubation, the mixture was applied to 293T cells (ATCC) ($1.2 \times 10^7$) in T175 flasks containing DMEM. The medium in these flasks was replaced 24 h post-transfection. At 48 h after transfection, the supernatant, which contained the virus-like particles (VLPs), was harvested, and subsequently passed through a 0.45µm syringe filter to ensure purity. For the luciferase assay, 100µl of this filtered supernatant was dispensed into each well of a 96-well plate, which already contained 30,000 MDMs per well. These cells were left to adhere overnight, allowing for the incorporation of the VLPs. The following day, the supernatant was removed, and the cells were washed with PBS (Corning). Then, 20 µL of cell lysis buffer (Promega) was added to each well, and the plate was gently rocked at room temperature for 15 minutes to facilitate cell lysis. The resulting lysate was then transferred to a white 96-well plate. To each well, 50 µL of reconstituted luciferase assay buffer was added and mixed thoroughly with the lysate. The luminescence generated by this reaction was measured immediately using EnSpire plate reader (PerkinElmer) to assess the efficiency of transfection and VLP incorporation.

Production of VSVΔG (G protein-deficient vesicular stomatitis virus) SARS-CoV-2 Spike pseudotyped virions and assessment of transduction efficiency by Renilla luciferase assay

For the preparation of virions, 293T cells were transfected with the spike plasmid, followed by inoculation with a generated working stock of VSVΔG-rLuc*G containing an integrated Renilla luciferase reporter gene to generate the pseudotyped VSVΔG-rLuc*SARS-CoV-2 (*51*). Pseudotyped virions were generated using Spike plasmids harboring mutations found in the WT

SARS-CoV-2 Spike (Wuhan-Hu-1). For the Renilla luciferase assay, 10µL of the virion stock was dispensed into each well of a 24-well plate, which already contained 100,000 Calu-3 cells per well. These cells were left to adhere overnight, allowing for the incorporation of the virions. The following day, the supernatant was removed, and the cells were washed with PBS (Corning). Then, 100 µL of cell lysis buffer (Promega) was added to each well, and the plate was gently rocked at room temperature for 15 minutes to facilitate cell lysis. The resulting lysate was then transferred to a white 96-well plate. To each well, 50 µL of reconstituted luciferase assay buffer was added and mixed thoroughly with the lysate. The luminescence generated by this reaction was measured immediately using EnSpire plate reader (PerkinElmer) to assess the efficiency of transduction and the pseudovirus incorporation.

**Production of SARS-CoV-2 replicon particles**

The pBAC SARS-CoV-2 Δ Spike WT plasmid (1 µg), was transfected into BHK-21 cells along with N and S expression vectors (0.5 µg each) in 24-well. The supernatant was replaced with fresh growth medium 16 hours post-transfection. The supernatant containing single-round infectious particles was collected and 0.45 µm-filtered 72 hours post-transfection. The supernatant was subsequently used to infect macrophages (in 96-well plate). To measure luciferase activity, an equal volume of supernatant from infected cells was mixed with Nano-Glo luciferase assay buffer and substrate and analyzed on an Infinite M Plex plate reader (Tecan).

**Isolation and Culture of Human AT2 Cells**

Human lungs from brain-dead donors who were declined for transplantation were obtained through Donor Network West for research and transported to UCSF under cold storage conditions. Human alveolar type II (AT2) cells were isolated from donor lungs as previously described (*53, 54*). Briefly, an uninjured lung lobe was selected based on gross inspection and chest imaging. The pulmonary vasculature was flushed, and distal airspaces were lavaged. Lung tissue was digested with elastase (13u/ml, Worthington Biochemical Corp.), minced, filtered, and subjected to density gradient centrifugation. $CD14^+$ cells were depleted, and AT2 cells were further enriched and purified by IgG panning.

Freshly isolated AT2 cells were seeded onto collagen I–coated Transwell inserts (0.4-µm pore size; Costar, Corning) in a 1:1 mixture of DMEM-H21 and Ham's F-12 supplemented with 10% fetal bovine serum (FBS) at a density of $1 \times 10^6$ cells per well. Cells reached confluence within 48 h. Medium in the apical compartment was removed daily to promote the formation of a tight epithelial monolayer. An air–liquid interface (ALI) was established 96–120 h after seeding, as indicated by the absence of fluid leakage from the basolateral compartment to the apical compartment. For coculture experiments, AT2 monolayers were washed with serum-free medium to remove residual serum prior to downstream experiments. MDMs were then seeded into the basolateral compartment of the Transwell system beneath the AT2 monolayer.

**Lung epithelial cell line**

Calu-6 epithelial cells (ATCC), genetically modified to stably express human angiotensin-converting enzyme 2 (hACE2; OriGene), were cultured in RPMI 1640 medium (Invitrogen), enriched with 10% fetal bovine serum (Gibco). For VLP transduction experiments, these cells were seeded onto 24-well plates at a density of $5 \times 10^4$ cells per well.

Calu-3 cells were maintained in high-glucose DMEM (Gibco) supplemented with 20% fetal bovine serum, 1% non-essential amino acids, 2 mM L-glutamine, 1 mM sodium pyruvate, 100 U/mL penicillin–streptomycin, and 1.5 g/L NaHCO3. For air–liquid interface (ALI) differentiation, Calu-3 cells were seeded onto 6-well Transwell inserts at a density of $5 \times 10^5$ cells per insert. Culture medium in both the apical and basolateral compartments was replaced every 2 days. On day 7 after seeding, medium was removed from the apical compartment to establish ALI conditions. Cells were maintained under ALI culture for an additional 11 days and used for experiments on day 18 after initial seeding.

**Primary human macrophages**

Primary human macrophages were generated from highly enriched peripheral monocytes. To obtain these cells, peripheral blood mononuclear cells (PBMCs) were first isolated from the blood of healthy donors (Vitalant) using Lymphoprep density gradient medium (StemCell Technologies). Subsequently, $CD14^+$ monocytes were selectively extracted from the PBMCs through negative selection, employing a combination of an antibody mixture designed for human monocyte enrichment and a magnetic column separation system (STEMCELL Technologies). Purified monocytes were cultured at a density of $1 \times 10^6$ cells/mL in serum-free Macrophage medium (StemCell Technologies), supplemented with M-CSF at a concentration of 50 ng/mL (PeproTech) for six days. For M1 polarization, monocyte-derived macrophages were stimulated with LPS (10 ng/mL) and IFN-γ (50 ng/mL) (both from PeproTech). For M2 polarization, cells were treated with IL-4 (10 ng/mL, PeproTech) following an 8-day differentiation protocol in the macrophage medium (STEMCELL Technologies).

**Inducing polarization in the THP-1 cell line**

The THP-1 cell line was maintained in RPMI 1640 medium (Invitrogen), supplemented with 10% fetal bovine serum (Gibco) and 2 mM L-glutamine (Invitrogen). To initiate differentiation into macrophage-like cells, the cells were treated with phorbol myristate acetate (PMA, Sigma-Aldrich) at a concentration of 150 nmol/L (PeproTech) for 24 h. Following this, the cells underwent multiple washes. To induce differentiation into the M1 phenotype, 20 ng/mL IFN-γ and 10 ng/mL LPS were added to the culture for 72 h. For M2 phenotype differentiation, the cells were treated with 20 ng/mL IL-4 and 20 ng/mL IL-13 for the same duration.

**Macrophage phenotyping**

The surface expression of CD14 (APC-Cy7), CD80 (PE), and CD206 (FITC) molecules (BD Biosciences) on macrophages was quantified using standard flow cytometry employing a BD LSR Fortessa X-20 instrument, with subsequent analysis carried out using FlowJo software (BD Biosciences). All macrophages were treated with Human BD Fc Block reagent (BD Biosciences) for 15 min before staining. Specific antibodies targeting these molecules, along with their respective isotype-matched control antibodies, were sourced from Caltag Laboratories.

**Transwell co-culture infection assay**

$3 \times 10^5$ human monocyte-derived macrophages were seeded onto Transwell plates using Macrophage medium (StemCell Technologies) enriched with M-CSF (50 ng/µL). Concurrently, an equal number of AT2 cells ($3 \times 10^5$) were cultured on the upper chamber of T-24 Transwell plate inserts (Corning) using DMEM high glucose 50%, F-12 50% mix medium (UCSF Media Production). The SARS-CoV-2 strain USA-WA1/2020 was then introduced to the cells at an MOI of 2. After two h, the media was replaced with fresh media. At 24, 48, and 72 h after infection, cell lysates were harvested using TRIzol (Invitrogen), and the culture supernatants were preserved at -80°C for further analysis by plaque assay.

**Reverse Transcription Quantitative PCR (RT-qPCR)**

Total cellular RNA was extracted utilizing TRIzol Reagent (Invitrogen) following the manufacturer's instructions. qRT–PCR was performed using 1 µg of total RNA with Luna Probe One-Step RT-qPCR 4× Mix with UDG (New England Biolabs) and TaqMan Gene Expression Assays (Thermo Fisher Scientific) according to the manufacturers' instructions. To validate low-abundance transcript detection, selected samples were additionally analyzed following pre-amplification using the TaqMan PreAmp Master Mix (Thermo Fisher Scientific). Quantitative PCR reactions were performed using a qTOWER3 thermocycler (Analytik Jena). For TMPRSS2, NRP1, CD16, cathepsin L, and TLR4 expression analyses, cDNA was synthesized using the iScript Reverse Transcription Supermix for RT-qPCR (BioRad), and qPCR was performed using Maxima SYBR Green/ROX qPCR Master Mix (Thermo Fisher Scientific) according to the manufacturers' instructions.

**Real-time quantification of phagocytosis**

MDMs were seeded at $1 \times 10^5$ cells per well in 96-well plates. Cells were stimulated with ORF8 (0.1 or 0.3 µg/mL) or SARS-CoV-2 nucleocapsid protein (0.3 µg/mL; GenScript) and incubated with pHrodo Green E. coli Bioparticles (100 µg/mL; Sartorius). Phagocytic activity was monitored in real time by quantifying the green fluorescence signal using an Incucyte SX5 live-cell analysis system.

**IL-10 ELISA**

ORF8 (Chempartner), at a concentration of 1 µg/mL, was added to the medium of MDM (M1) macrophages. After 48 h, LPS (PeproTech) at concentrations of 1, 10, or 100 ng/mL was introduced. Following 24 h, the medium was collected. This medium was then applied to anti-human IL-10 pre-coated 96-well plates, and an ELISA was conducted as per the manufacturer's instructions (BioGems). The IL-10 concentration was determined using the standard curve.

**FLICA assay**

MDMs were cultured with VLPs alone or with VLPs supplemented with recombinant ORF8 (1 μg/mL). Activated caspase-1 was detected using either green fluorescent FAM-FLICA or far-red fluorescent FLICA 660 reagents (ImmunoChemistry Technologies) according to the manufacturer's instructions. After 30 min of incubation, cells were washed according to the manufacturer's protocol and stained with Zombie Violet viability dye (BioLegend). Samples were subsequently analyzed using a BD LSRFortessa X-20 flow cytometer (BD Biosciences).

**NF-κB-/AP-1-reporter assay**

The reporter constructs were generated using the pSIRV transfer vector together with Moloney murine leukemia virus (MMLV) gag/pol (Addgene) and VSV-G packaging plasmids. NF-κB- or AP-1-responsive promoters were used to drive secreted NanoLuc luciferase expression as a readout for NF-κB and AP-1 signaling activation, respectively. These retroviral vectors were transduced into MDMs, followed by stimulation with ORF8 or IL-17 4 days later. At the time of sample collection, culture supernatants were completely removed, cells were washed three times with PBS, and fresh medium was added prior to downstream analysis.

**Proteomics analysis**

Abundance proteomics sample preparation. Each of the 36 frozen cell pellets (sample details in Table S4) was resuspended in 100μL of urea lysis buffer (8M urea, 150mM NaCl2, 100mM Tris) on ice, sonicated for 30 seconds, prior to one round of freeze thaw. Protein concentration of the resulting lysates was determined by Bradford assay. For each lysate, 50μg was reduced with 2mM tris(2-carboxyethyl)phosphine (Sigma, C4706) for 30 minutes at room temperature and alkylated with iodoacetamide final concentration 10mM for 30 minutes at room temperature in the dark. Iodoacetamide was quenched by addition of DTT at final concentration 10mM and incubation in the dark at room temperature for 30 minutes. Buffer exchange and protein digestion were performed on a KingFisher Flex unit using a protein aggregation approach. In this digestion protocol the setup is as follows: plate #1 stores the 96-well comb; plate #2 contains the lysate/bead mixtures; plate #3-5 contain 95% acetonitrile wash solutions; plates #6-7 contain 70% ethanol wash solutions; and plate #8 contains the digestion elution buffer. Prior to starting the automated runs, the lysate/bead mixtures are prepared by mixing together 20μL MagReSyn Amine beads (Resyn Biosciences) with 100μL of lysate and 280μL of 100% acetonitrile (to plate #2). Protein aggregation on beads was induced by 4 cycles of mixing (1 min), pause (5 min), and collection of the magnetic beads (24 min total time). Beads with bound protein were then transferred to plate #3 and released into 150 μL of 95% acetonitrile and washed by 5 cycles of mixing. Washing cycle is repeated two times in plates #4 and #5. Beads are then transferred to plate #6 and released into 150 µL of 70% ethanol and washed by 5 cycles of mixing. This cycle was repeated once more in plate #7. Finally, protein bound beads were released into plate #8 with freshly prepared digestion elution buffer (150 µL of 50 mM ammonium bicarbonate (pH 7.8) with 0.5µg Lys-C and 0.5µg trypsin). Plate #8 with the digesting proteins was then sealed, and proteolytic digestion proceeded overnight at 37 °C with agitation at 800 RPM (Eppendorf ThermoMixer). The resulting peptides were then filtered by 0.45 µm membranes, acidified with 5µL formic acid, and then dried by vacuum centrifugation.

Protein abundance mass spectrometry (MS) data acquisition and analysis. For MS acquisition, dried samples were resuspended in 25µL of sample buffer (0.1% formic acid in water) and 1µL of peptide was injected on a Orbitrap Exploris 480 mass spectrometer (Thermo Fisher Scientific) coupled to a Vanquish Neo liquid chromatography instrument (Thermo Fisher Scientific). Briefly, peptides were separated on a 150µm x 15cm PepSep column (Bruker, 1.5µm beads) over a 90min gradient at a flow rate of 600 nL/minute as described in Table S5. Buffer A consisted of 0.1% formic acid (FA) in water, and buffer B was 0.1% FA in 80% acetonitrile. Spectra were continuously acquired in a data-dependent acquisition (DDA) mode. One full scan was acquired in the Orbitrap (350-1250 m/z at 120,000 resolution with a normalized AGC target of 300%) followed by as many MS/MS scans as could be acquired on the most abundant ions in 2 seconds in the Orbitrap (1500 resolution; Normalized collision energy type; HCD collision energy of 28%; normalized AGC target of 200 %, maximum injection time set to Auto, and isolation window of 1.6 m/z). Singly and unassigned charge states were rejected, and dynamic exclusion was enabled after n=1 time, with an exclusion duration of 20 seconds (tolerance of ±10 ppm). Detailed MS acquisition parameters are reported in Table S6. Raw MS files were analyzed using Fragpipe (version 21.1). MS/MS spectra were searched against the human proteome (UniProt reviewed, downloaded 09 May 2024; concatenated with equal sized reversed database and common contaminants) using default parameters (Table S7).

Peptide ion intensities from the output of Fragpipe (Table S7) were summarized to protein intensities using the R Bioconductor package MSstats (version 4.8.3). The functions dataProcess and groupComparison were applied with default settings, except for setting "MBimpute = FALSE" in dataProcess. Proteins with an adjusted p-value ≤ 0.05 and an absolute $\log_2$ fold change > 1 were considered significant. Significantly changed genes were tested for Gene Ontology (GO) term enrichment, including Biological Process (BP), Molecular Function (MF), and Cellular Component (CC), using the enricher function from the clusterProfiler package (version 4.8.1). GO annotations were retrieved using clusterProfiler:get_GO_data. Enriched GO terms with adjusted p-values < 0.1 were considered significant. To remove redundancy among GO terms, we constructed a GO term tree based on pairwise distances defined as 1 - Jaccard similarity coefficient of shared genes. The tree was cut at a height of h = 0.99 to identify clusters of similar terms. Within each cluster, the representative term was selected by choosing the broadest significant GO term.

**Bulk RNA sequencing**

Monocytes (1 × $10^6$ cells) were seeded in 6-well plates and differentiated into macrophages. Cells were stimulated with ORF8 (1 µg/mL) for 3 h prior to analysis. Cell lysates were then prepared in 150 µL of DNA/RNA Shield (Zymo Research). Bulk RNA sequencing was performed by Plasmidsaurus. RNA quality was assessed prior to library preparation, and sequencing libraries were generated according to the manufacturer's standard protocols. Libraries were sequenced on an Illumina platform. Reads were aligned to the human reference genome (GRCh38), and differential expression analysis was performed using DESeq2 with an adjusted p value cutoff of 0.05.

**C57BL/6 and BALB/c wild-type mouse SARS-CoV-2 infection model**

All protocols concerning animal use were approved (AN203103-00F) by the Institutional Animal Care and Use committees at the University of California, San Francisco and Gladstone Institutes and conducted in strict accordance with the National Institutes of Health Guide for the Care and Use of Laboratory Animal. C57BL/6J mice were obtained from The Jackson Laboratory, and BALB/c mice were obtained from Charles River Laboratories. Mice were housed in a temperature- and humidity-controlled specific pathogen-free facility under a 12 h light/dark cycle with ad libitum access to water and standard laboratory chow. We constructed a mouse-adapted (MA)-SARS-CoV-2 (Spike: Q498Y/P499T54) using pGLUE42. MA-SARS-CoV-2 (1 × $10^3$ PFU in 40 µL) was administered intranasally under anesthesia to 6–8-week-old mice. For MA-SARS-CoV-2 StopORF8 experiments, purified ORF8 protein (10 µg; ChemPartner) was administered intraperitoneally at 24 and 48 h post-vaccination. In prevention assays, wild-type mice received intranasal delivery of the peptide (25 µg in 40 µL), followed 30 min later by intranasal inoculation with MA-SARS-CoV-2 (1 × $10^3$ PFU in 40 µL). Mice were euthanized 24 h after peptide treatment. For intervention studies, IL17RA antibody (100 µg; R&D Systems) was administered intraperitoneally 2 h after ORF8 injection, and mice were euthanized 72 h after viral challenge. Mouse IgG1 isotype control (R&D Systems) was used as a negative control. For assessment of lung fibrosis, mice were euthanized 5 days after infection and lungs were collected for analysis.

F4/$80^+$ macrophages and EpCAM$^+$ epithelial cells were isolated from lungs 2 days after viral inoculation. Excised lungs were minced with scissors and digested in RPMI containing DNase I (100 µg/mL; Roche) and Liberase TL (10 µg/mL; Sigma-Aldrich), followed by mechanical dissociation using a gentleMACS Octo Dissociator (Miltenyi Biotec). Cell suspensions were separated with Anti-F4/80 MicroBeads and CD326 (EpCAM) MicroBeads (Miltenyi Biotec) and subjected to RT–qPCR analysis.

**Histology**

Mouse lung tissues were fixed in 4% PFA (Sigma Aldrich) for 24 hours, washed three times with PBS and stored in 70% ethanol. All the stainings were performed at Histo-Tec Laboratory, Inc (Hayward, CA). Samples were processed, embedded in paraffin, and sectioned at 4µm. The slides were dewaxed using xylene and alcohol-based dewaxing solutions. Epitope retrieval was performed by heat-induced epitope retrieval of the formalin-fixed, paraffin-embedded tissue using 10mM citrate-based pH 6 solution for 20 mins at 95°C. The tissues were stained for H&E and Caspase-1 (Abcam) and dried, coverslipped, and visualized using Axioscan 7 (ZEISS) at 10X and 40X. For Masson's trichrome staining, nuclei were stained with Weigert's iron hematoxylin solution, followed by staining with Biebrich scarlet-acid fuchsin solution to visualize cytoplasm and muscle fibers. Sections were then treated with phosphomolybdic-phosphotungstic acid solution and subsequently stained with aniline blue to label collagen fibers. Finally, slides were differenciated in 1% acetic acid, dehydrated in graded ethanol, cleared in xylene, and mounted with a resinous medium. All slides were interpreted by a board-certified pathologist. Caspase-1-positive cells and pulmonary fibrotic area were quantified in Fiji/ImageJ.

**Fig. S1.**

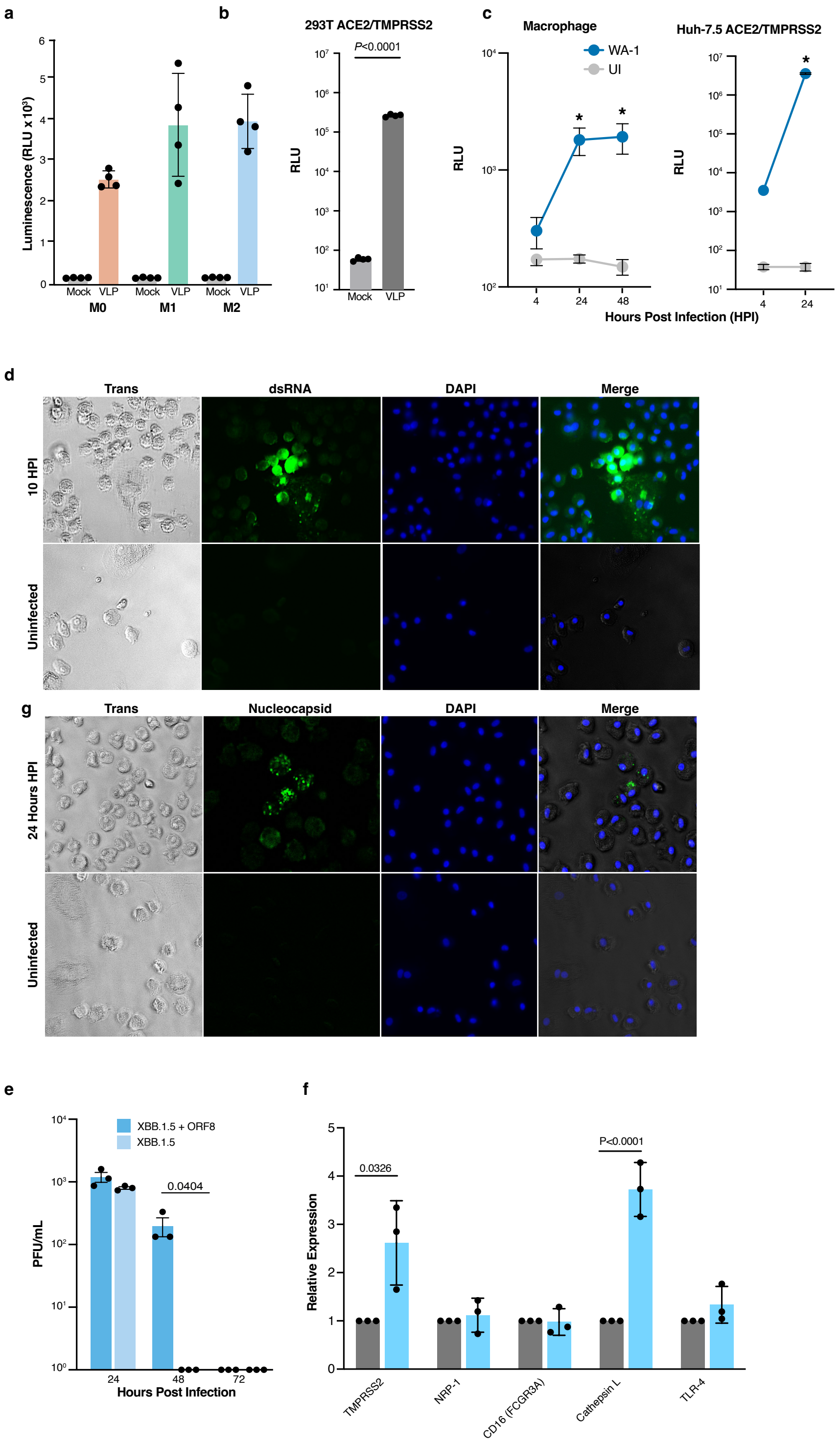


**Figure S1: MDMs support limited and transient production of infectious SARS-CoV-2.** (A) MDMs (M0, M1, and M2) were exposed to luciferase-encoding VLPs for 24 h, and luciferase activity was subsequently quantified. Data are mean ± SD. (B) 293T-ACE2/TMPRSS2

cells were exposed to luciferase-encoding VLPs for 24 h, followed by quantification of luciferase activity. (C) MDMs and Huh-7.5 ACE2/TMPRSS2 cells were exposed to replicons, and NanoLuc activity in the supernatant was quantified as a measure of SARS-CoV-2 RNA replication. *$p < 0.05$. (D) MDMs were infected with replication-competent SARS-CoV-2 at MOI 1; dsRNA was detected at 10 h (Top) followed by N protein staining at 24 h (Bottom). (E) MDMs were infected with SARS-CoV-2 XBB.1.5 or XBB.1.5 supplemented with extracellular ORF8 at an MOI of 2. Culture supernatants were collected at 24-, 48-, and 72-h post-infection, and viral titers were quantified by plaque assay. Statistical significance was determined using two-tailed Welch's t-test. (F) MDMs were treated with ORF8 (1 µg/mL). Expression of the indicated genes was quantified by qPCR normalized to HPRT. Data are presented as fold change relative to unstimulated ± SD of three independent experiments. Statistical significance was determined using two-tailed Welch's t-test.

Fig.S2.

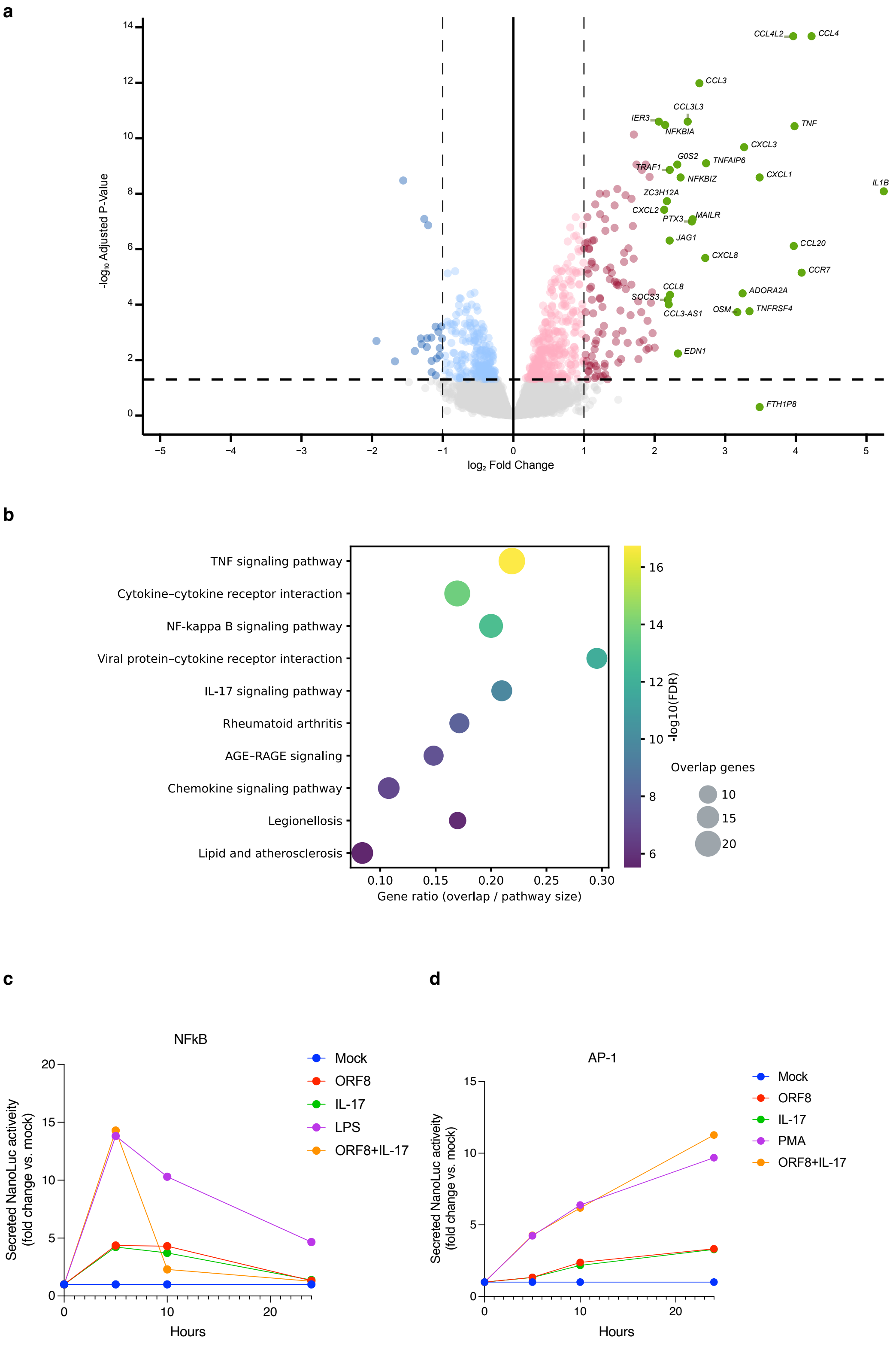

a
CCL4L2
CCL4
CCL3
CCL3L3
IER3
NFKBIA
TNF
CXCL3
G0S2
TNFAIP6
TRAF1
NFKBIZ
CXCL1
IL1B
ZC3H12A
CXCL2
PTX3
MAILR
JAG1
CCL20
CXCL8
CCR7
CCL8
ADORA2A
SOCS3
CCL3-AS1
OSM
TNFRSF4
EDN1
FTH1P8
-log10 Adjusted P-Value
log2 Fold Change
b
TNF signaling pathway
Cytokine–cytokine receptor interaction
NF-kappa B signaling pathway
Viral protein–cytokine receptor interaction
IL-17 signaling pathway
Rheumatoid arthritis
AGE–RAGE signaling
Chemokine signaling pathway
Legionellosis
Lipid and atherosclerosis
Gene ratio (overlap / pathway size)
-log10(FDR)
Overlap genes
c
NFkB
Secreted NanoLuc acteivity (fold change vs. mock)
Hours
Mock
ORF8
IL-17
LPS
ORF8+IL-17
d
AP-1
Secreted NanoLuc acteivity (fold change vs. mock)
Hours
Mock
ORF8
IL-17
PMA
ORF8+IL-17

**Figure S2: ORF8-Induced NF-κB/AP-1 Signaling Responses in MDMs.**

**(A)** MDMs were stimulated with ORF8 (1 μg/mL) for 3 h, followed by RNA-seq analysis in biological triplicates and comparison with mock-treated controls. Red dots indicate upregulated genes, blue dots indicate downregulated genes, and green dots indicate the top 30 upregulated genes. **(B)** KEGG pathway enrichment analysis of significantly upregulated genes (adjusted $P < 0.05$, log2 fold change $> 1$) is shown as a dot plot. Dot size represents gene ratio, and color indicates false discovery rate (FDR). **(C and D)** MDMs transduced with NF-κB (C) or AP-1 (D) NanoLuc reporter lentiviruses were treated with ORF8 (1 μg/mL), IL-17 (200 ng/mL), LPS (100 ng/mL), or ORF8 plus IL-17. Culture supernatants were collected at 5, 10, and 24 h post-treatment, and NanoLuc activity was quantified. At the 5- and 10-h time points, supernatants were completely removed, cells were washed with PBS, and fresh medium containing the indicated stimuli was added. Experiments were performed in biological triplicates, and data are presented as mean values.

**Fig.S3.**

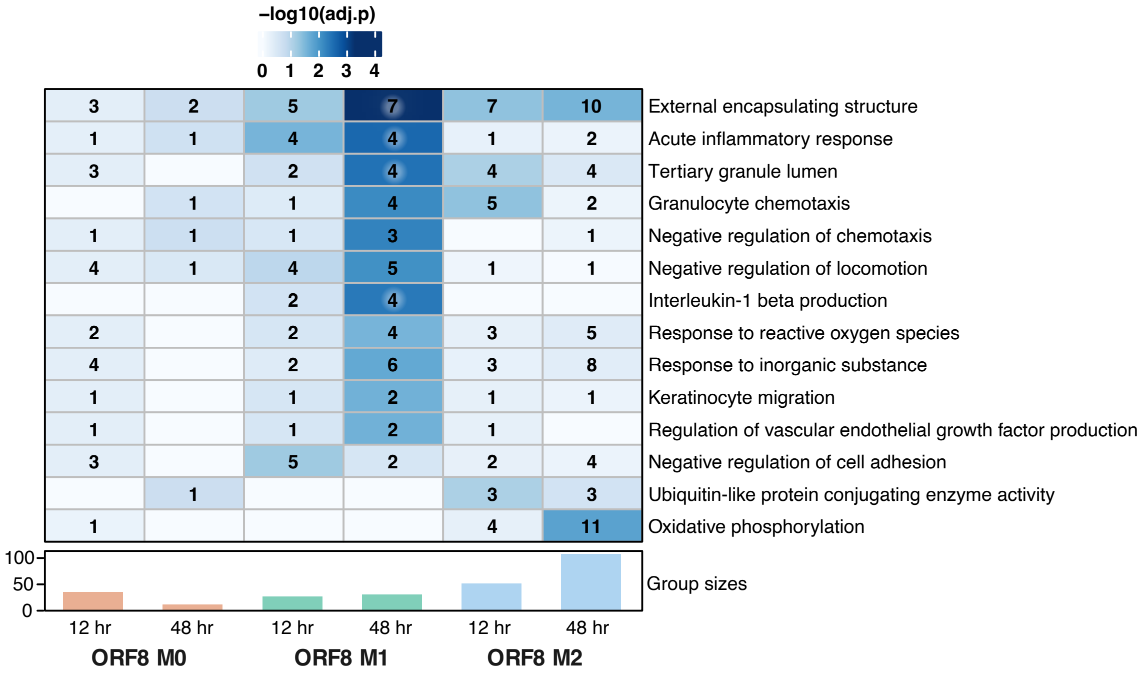


**Figure S3: GO enrichment analysis of ORF8-exposed macrophages.** Gene Ontology (GO) term enrichment analysis was performed on differentially expressed proteins identified by proteomic profiling of M0, M1, and M2 macrophages following exposure to extracellular ORF8. Significantly enriched biological processes, molecular functions, and cellular components are shown, with enrichment scores calculated relative to control condition.

**Fig. S4.**

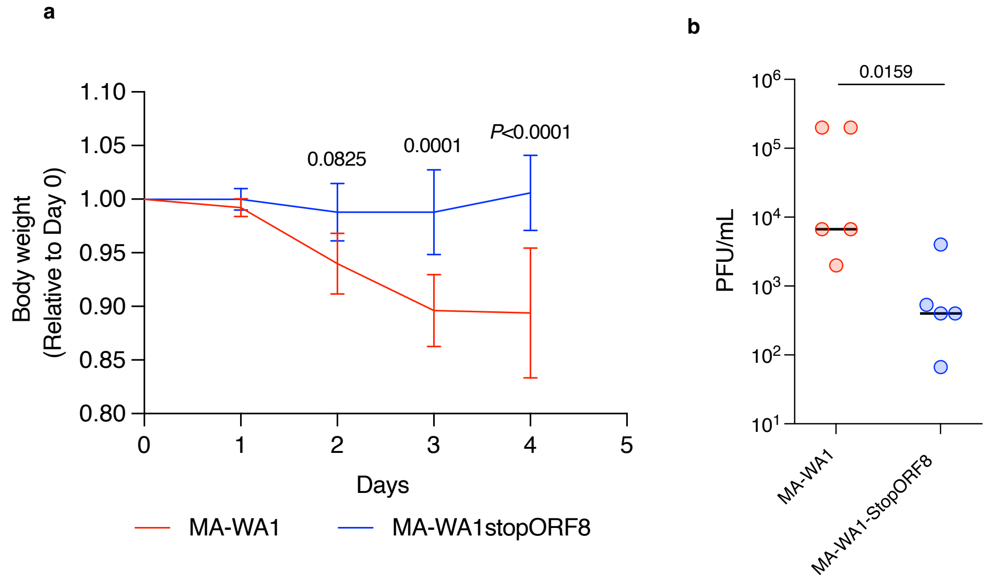


**Figure S4: ORF8 Determines Disease Outcome in Aged Mice.**

(A and B) One-year-old male BALB/c mice were intranasally infected with mouse-adapted WA-1 (n = 5) or mouse-adapted WA-1-StopORF8 (n = 5). Body weight was monitored daily (A), and lungs were harvested at 4 days post-infection for plaque assay analysis (B). Statistical significance in (A) was determined by two-way ANOVA with Šídák’s multiple-comparisons test, whereas (B) was analyzed using a two-tailed Welch’s t-test.

**Fig. S5.**

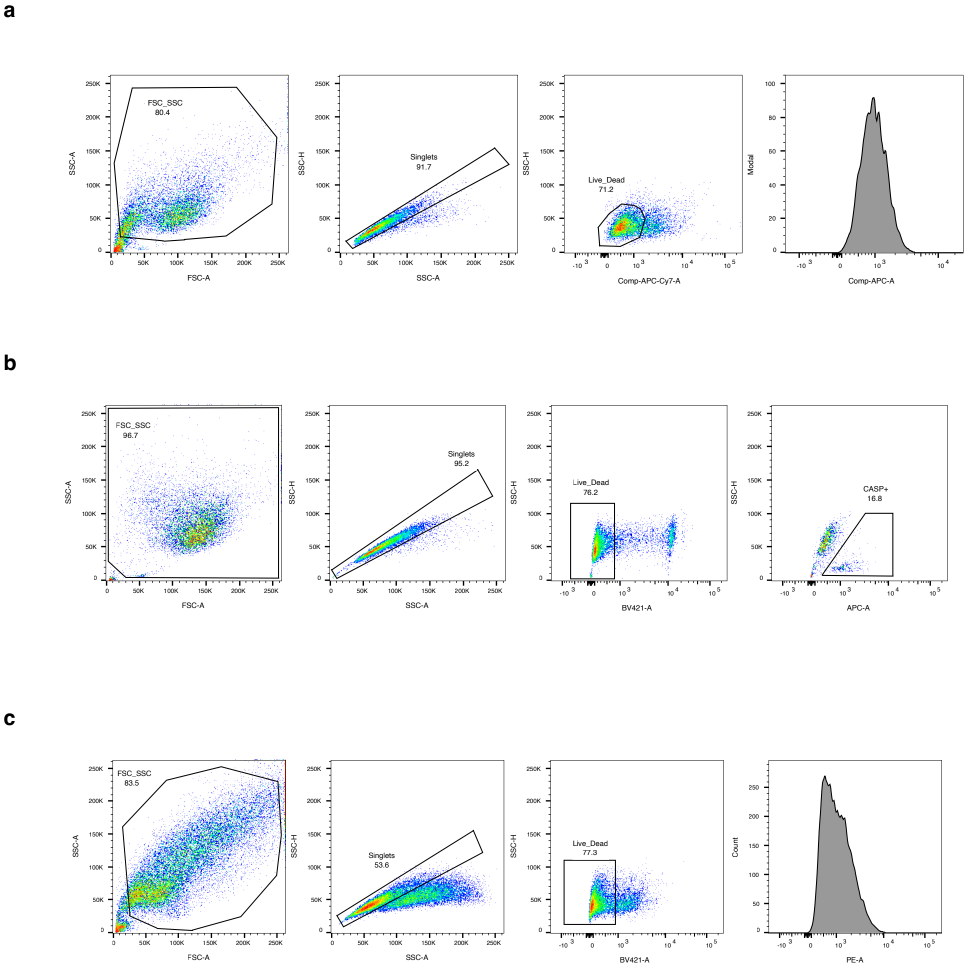


**Figure S5: Flow cytometry gating strategy.**

(A) Gating strategy for ACE2 expression. (B) Gating strategy for FLICA analysis. (C) Gating strategy for TMPRSS2 expression

**Table S1**: MSstats quantification of protein lysates from M0, M1, and M2 primary monocyte derived macrophage (MDM) exposed to ORF8 for 12 and 48 hours

**Table S2**: Gene Ontology (GO) enrichment analysis of proteins identified in M0, M1, and M2 primary MDMs exposed to ORF8 for 12 and 48 hours

**Table S3**: qPCR primer sequences

**Table S4**: Metadata overview for the proteomic abundance dataset from M0, M1, and M2 primary MDMs exposed to ORF8

**Table S5**: Liquid Chromatography (LC) parameters used for collection of proteomic abundance data

**Table S6**: Orbitrap Exploris 480 mass spectrometry (MS) parameters used for collection of proteomic abundance data

**Table S7**: Fragpipe parameters used for searching proteomic data
